# Gradual decrease in the inter-population diversity of HIV-1 Env among group M lineages worldwide

**DOI:** 10.1101/753285

**Authors:** Changze Han, Jacklyn Johnson, Rentian Dong, Raghavendranath Kandula, Alexa J. Kort, Maria Wong, Tianbao Yang, Patrick J. Breheny, Grant D. Brown, Hillel Haim

## Abstract

HIV-1 group M was transmitted to humans nearly one century ago. The virus has since diversified to form distinct clades, which spread to multiple regions worldwide. Of the different proteins encoded by HIV-1, the envelope glycoproteins (Envs) have diversified most rapidly in all infected populations. We compared the range of variants that emerged during the AIDS pandemic in diverse HIV-1 clades and distinct geographic regions. Our analyses focused on two components of Env that contain multiple epitopes of broadly-neutralizing antibodies: the glycan shield and apex domain. Interestingly, at each Env position, the amino acid in the inferred clade ancestor was replaced by a unique combination of emerging variants. Key antigenic sites and genetic signatures of vaccine protection have gradually evolved toward conserved frequency distributions (FDs) of all amino acids. FDs are specific for position and clade and are highly conserved in populations from different regions. Remarkably, founder effects of Env mutations in distinct clades and recently-infected regions were significantly reduced during the epidemic by evolution of each site toward the position-specific FD. These findings suggest that the selective pressures that guide evolution of Env are conserved in different populations. They are sufficiently strong to reduce founder effects at the clade and regional levels and have significantly altered the distribution of Env forms that circulate worldwide. Consequently, the intra-population diversity of the Env protein continues to increase whereas the inter-population diversity is gradually decreasing.

**Importance:** The Env protein of HIV-1 is the primary target in AIDS vaccine design. Due to frequent mutations, new Env variants continuously emerge in the population. The increasing number of Env forms and apparent randomness of the changes limit our ability to design broadly-effective vaccines. We examined the populations-level changes that occurred in Env during the AIDS epidemic. Each position of the molecule has evolved toward a specific combination of amino acids. Similar changes occurred in different HIV-1 subtypes and geographic regions toward the same sets of forms, often from distinct ancestral sequences. Such conserved patterns of evolution define a new framework for designing vaccines that are tailored to the unique combination of Env variants expected to circulate in each population.

## Introduction

The founder virus of HIV-1 group M was transmitted to humans in the early part of the 20^th^ century (1–3). Due to low fidelity of the viral replication machinery, frequent mutations are introduced in the HIV-1 genome (1, 2). The group M founder thus gradually diversified to create different genetic lineages (clades), which spread to multiple regions of the world (3, 4). In some regions, a single founder was introduced that accounts for most circulating strains, such as the clade B lineage in Korea (5) or the clade C lineage in India (6, 7). In other regions, multiple founder lineages were introduced. The envelope glycoprotein (Env) on the surface of the virus is the most diverse of all proteins encoded by HIV-1. More than 20% of Env residues can differ among viruses from the same clade (8–11). This protein continues to diversify at a population level (11–14). Many epitopes recognized by broadly-neutralizing antibodies (BNAbs) were strictly conserved during the early years of the epidemic and are now present in a significantly smaller proportion of circulating strains (12). The antigenic diversity of Env poses a major challenge to the ability of vaccines to elicit a broadly-effective antibody response (15–17). Several studies have examined the within-population changes that occurred in Env during the pandemic (11–14). However, less is known about the between-population changes. Are the same variants of Env emerging in diverse clades and distinct geographic regions? Is Env evolving toward preferred structural forms? Are founder effects of the virus in diverse clades and newly-established regions stable over the course of time?

To address these questions, we examined the population-level changes that occurred in the Env protein during the AIDS pandemic. We focused on two components of Env that contain multiple BNAb epitopes: (i) the glycan shield of gp120, composed of multiple N-linked glycosylation sites that adorn the surface of the molecule (18, 19), and (ii) the second variable loop (V2) segment at the trimer apex, which also contains two signatures of vaccine protection identified in sieve analysis of the RV144 trial results (20–23). Significant population-level changes have occurred during the AIDS pandemic in the glycan shield and V2 apex. Interestingly, each position of Env has evolved toward a unique combination (frequency distribution, FD) of amino acids that is highly conserved in different populations worldwide. FDs also exhibit clear clade-specific patterns; they are conserved in distinct geographic regions and in monophyletic and paraphyletic subclade groups. Therefore, our findings suggest that the same selective pressures guide evolution of each position of Env in different populations. Such pressures are sufficiently strong to reduce founder effects of the virus at the clade and regional levels and direct each site to the same unique combinations of variants.

## Results

### Population-level changes in the glycan shield of Env are unique for each position and follow similar patterns in distinct regions worldwide

To understand the pressures applied on Env at a population level, we compared the historical changes in amino acid sequence at different sites on the trimer in diverse clades and geographic regions. We first examined six adjacent positions on the glycan shield of Env (**Fig. 1A**). All six sites were occupied by a potential N-linked glycosylation site (PNGS) in the group M ancestor and are located at positions generally unaffected by insertions or deletions. Glycans at these sites play critical roles in protecting Env from targeting antibodies and also serve as conserved targets of BNAbs (24–27). Historical changes in PNGS frequency at these sites were calculated among isolates from clades B, C, A1 and CRF01_AE, using 1942, 1248, 335 and 543 unique-patient sequences, respectively (see phylogenetic trees of all Env panels in **Fig. S1** and amino acid alignments in **Datasets S1-S4**). Three of the six sites (289, 295 and 332) showed clear clade-specific patterns (**Fig. 1B**). Clades that contained a PNGS in their ancestral sequence maintained conserved frequencies during the pandemic. Clades that did not contain an ancestral PNGS often showed an increase in frequency of this motif, although the level remained distinct from other clades. Such clade-specific patterns were less prominent for the sites at positions 339, 386 and 392 (**Fig. 1C**). Interestingly, the frequency of PNGSs at position 339 in CRF01_AE rapidly increased during the pandemic to replace the ancestral Asn (not part of a PNGS motif).

**Fig. 1.**
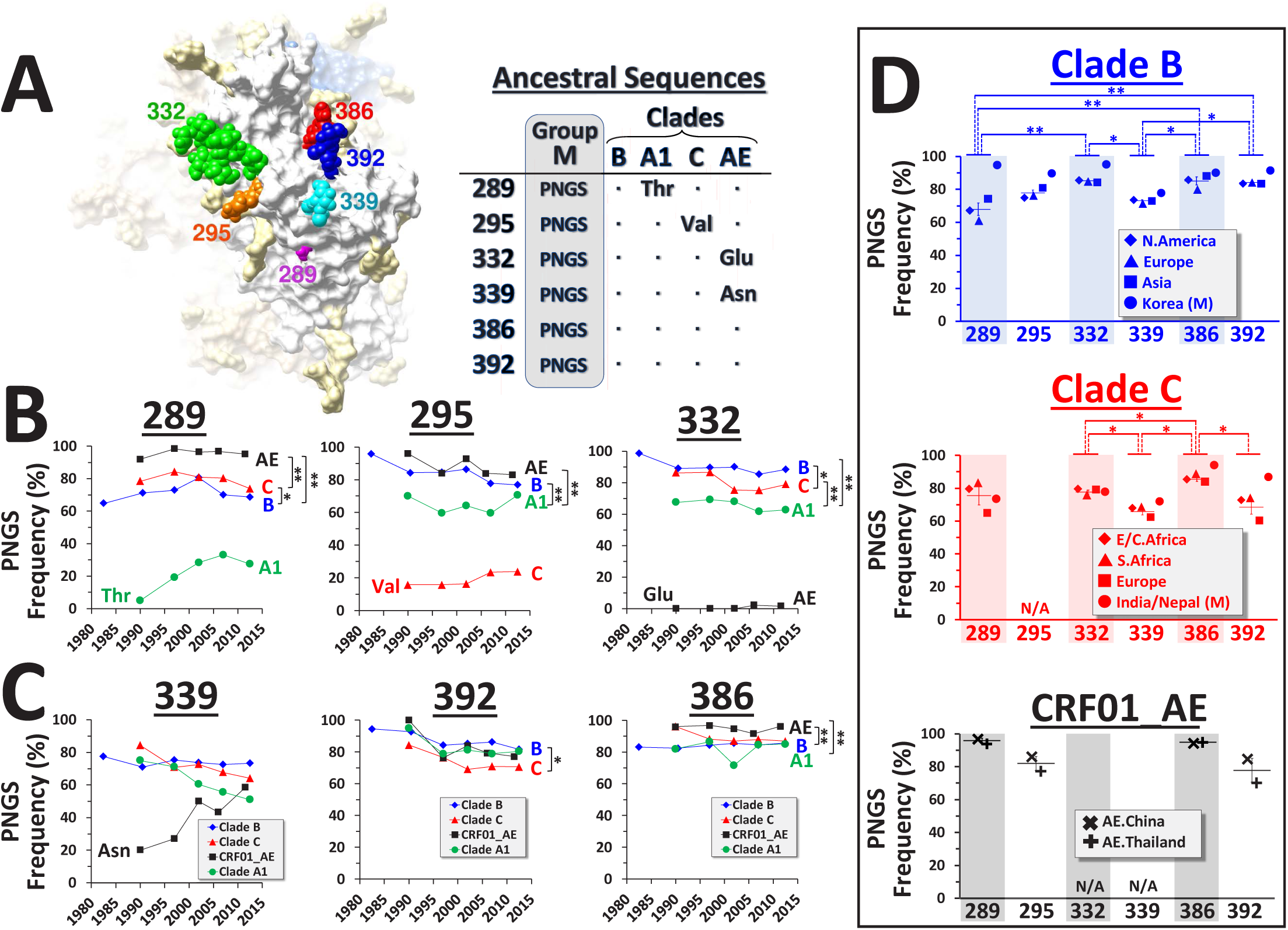
Population-level frequencies of PNGSs at six positions of gp120 are clade-specific and conserved in different geographic regions. **(A)** Cryo-EM image of the BG505 SOSIP.664 Env trimer (PDB ID 4TVP). Six Env positions that contain a PNGS motif in the inferred group M ancestor are shown. Position 289 is occupied by a Thr residue in the clade A1-derived BG505 Env. Ancestral sequences at these positions in four group M clades are shown to the right. **(B,C)** Historical changes in PNGS frequencies at the six positions of gp120. Envs were isolated from samples collected worldwide between 1979 and 2015 (one Env per patient). PNGS frequencies are calculated for consecutive 5-7-year periods in clades B (1,942 patients, blue), C (1,248 patients, red), A1 (335 patients, green) and CRF01_AE (543 patients, black). This color-coding scheme is applied throughout the manuscript. The residue found in the inferred ancestor of each clade (if not a PNGS) is indicated. P values for a one-way ANOVA test that compares all time points between clades that contain a PNGS in their ancestral sequence: *, P<0.05; **, P<0.01. **(D)** PNGS frequencies in the indicated regions among recently-circulating strains (see year ranges and country compositions in **Table S1**). Averages and position specificity of the patterns (using a one-way ANOVA test) were calculated among the regional panels that compose the paraphyletic groups of clades B and C: *, P<0.05; **, P<0.01. “M” denotes Envs from the monophyletic clusters that circulate in Korea and India/Nepal. N/A, position is not occupied by a PNGS in the clade ancestor. Error bars, standard errors of the mean (SEM).

To examine position-specificity of these patterns, we compared PNGS frequencies in populations from distinct geographic regions (**Fig. 1D**). Envs isolated from recently-collected samples were analyzed, defined for most regional panels as the year range 2007-2015 (see composition of all panels in **Table S1**). PNGS frequencies were compared between populations that contained this motif in their inferred regional founders (for the monophyletic lineages) or their clade-ancestral sequences (for the paraphyletic groups). Interestingly, PNGS frequencies were unique for each position and conserved in the different regions. The range of these values was surprisingly narrow for the regional panels in the paraphyletic group (from North America, Europe and Asia); 84 to 85 percent at position 332, 73 to 74 percent at position 339 and 83 to 84 at position 392 (see position specificity of the patterns in **Fig, 1D** and combined data for all four clades in **Fig. S2**). The monophyletic group from Korea was introduced into this region in the late 1980’s or early 1990’s (5) (see tree in **Fig. S1A**). Consequently, the frequency of PNGSs at all sites was higher in the Korean lineage, although similar position-specific patterns were still observed. Clade C also showed position-specific frequencies, which were similar in the monophyletic lineage from India and Nepal (6, 7) and regional panels of the paraphyletic group from S. Africa, Eastern/Central (E/C) Africa and Europe (see **Table S1**).

Therefore, the frequency of PNGSs at six adjacent positions of Env that contained this motif in their clade ancestors has changed during the pandemic in a position- and clade-specific manner. PNGS frequencies among currently-circulating strains occupy narrow ranges of values, which are conserved in different populations worldwide.

### Population-level frequencies of emerging variants are unique for each position and conserved in different regions worldwide

To further investigate these patterns, we examined changes in frequency of all residues that emerged to replace the ancestral PNGS. The frequency distribution (FD) pattern of emerging residues seemed distinct for each site (compare 392, 386 and 339 in **Fig. 2A**, all six positions in **Fig. S3, A-D**, and historical patterns in **Fig. S4**). At some positions (e.g., 392), similar FDs were observed in the diverse clades; frequencies as low as 2% were conserved. At other positions (e.g., 339), more variation was observed between clades. Comparison of residue frequencies in the regional panels of each clade showed clear position-specific patterns (see Ser and Asp in **Fig. 2, B and C**, and other residues in **Fig. S5**). Frequencies in the clade B groups from N. America, Europe and Korea occupied narrow ranges of values, even at low frequencies. Clade C also showed conserved frequencies of residues in Europe, S. Africa and E/C Africa. A similar profile was observed for the monophyletic clade C lineage from India and Nepal (**Fig. S3D).**

**Fig. 2.**
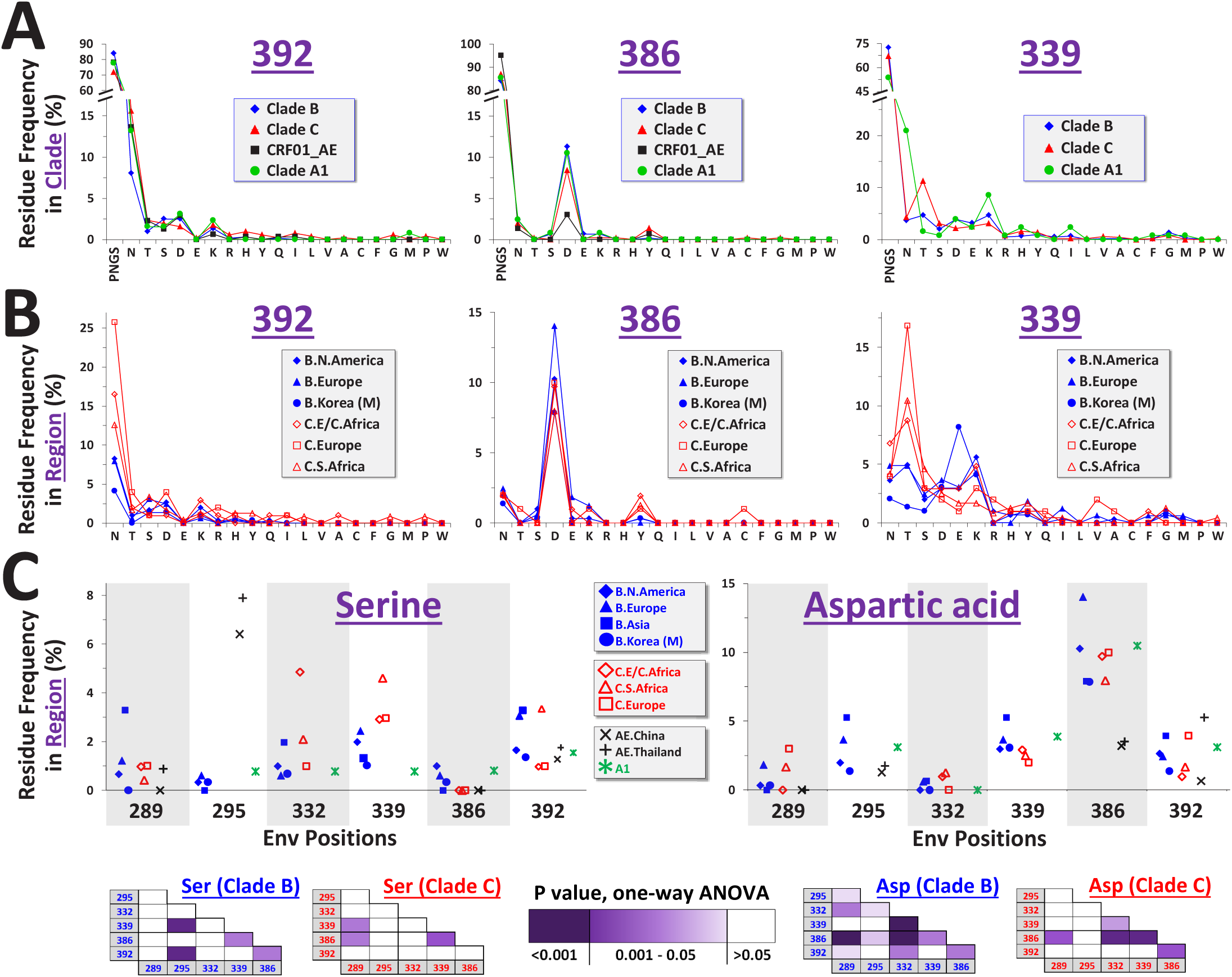
Frequencies of residues that replaced the clade-ancestral PNGS motif are position specific and occupy narrow ranges of values. **(A)** Frequency distributions (FDs) of residues at positions 392, 386 and 339 in clades B, C, A1 and CRF01-AE, calculated among recently-circulating strains. Clades that contained a PNGS motif in their ancestral sequence at each position are shown. Residues are labeled by single-letter code. “N” indicates Asn that is not part of a PNGS motif. Profiles for all six positions are found in **Fig. S3A**. **(B)** Positional FDs in the indicated regional panels among recently-circulating strains (see also **Fig. S3**, **B** and **C**). **(C)** Frequencies of Ser and Asp in regional panels of clades B, C, A1 and CRF01_AE. A one-way ANOVA test that compares residue frequencies at the different positions was performed for clades B and C; cells are color-coded by P values as indicated.

**Fig. 3.**
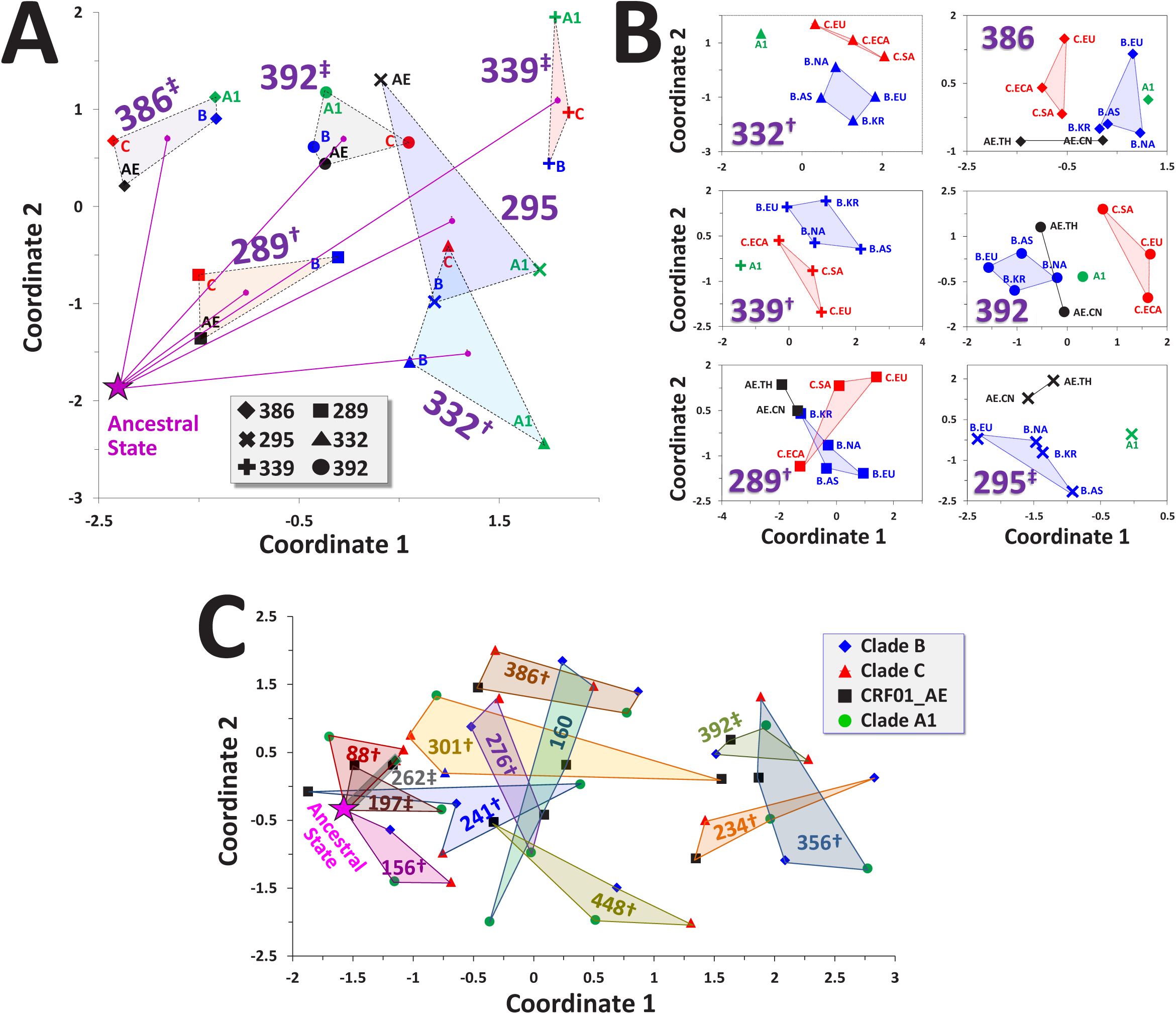
Frequency distributions of residues that replaced the ancestral PNGS motif are specific for each Env position and HIV-1 clade. **(A)** Relationships between positional FDs in diverse clades. Positional FD profiles of clades that contained a PNGS in their inferred ancestral sequence are compared. Each data point represents the FD at the indicated position in a given clade. Location of a profile composed solely of PNGSs is labeled “Ancestral State”. Dashed lines connect FDs for the same position, and a line is drawn from the ancestral state to the centroid of each. Position-specificity of the patterns was calculated by a permutation-based test: †, P<0.05; ‡, P<0.005. **(B)** Clade specificity of FDs. Positional FDs were calculated for the indicated regions among recently-circulating strains. Only regional panels that contained a PNGS in their ancestral/founder sequence are shown. Clade specificity of the patterns is indicated: †, P<0.05; ‡, P<0.005. **(C)** Relationships between FDs of 13 positions of gp120 that contain a PNGS in the inferred ancestral sequence of all four clades. Data points represent FDs at the indicated positions calculated among recently-circulating strains. FDs for the same position are connected by solid lines. Position-specificity of the patterns is indicated: †, P<0.05; ‡, P<0.005.

Therefore, distinct and conserved profiles of variants have emerged at each Env position to replace the ancestral PNGS. These specificity patterns for position and clade cannot be attributed to differential synonymous codon usage in the clade-ancestral sequences (**Supp Fig. S3E**). Conserved patterns among panels that compose the paraphyletic groups of each clade could potentially result from ‘mixing’ of the viruses between regions. However, the similar profiles observed in the monophyletic lineages suggest that the patterns likely result from conserved selective pressures applied in the different populations.

### Frequency distribution profiles are specific for each Env position and HIV-1 clade

To determine specificity of the complete profiles (composed of all emerging residues) we examined the relationships between FDs at the different positions in diverse clades and geographic regions. For this purpose, the positional FD in each population is treated as a 21-feature vector that describes the log_10_ frequency of all 20 residues and a PNGS. Euclidean distances between vectors are calculated as a measure of differences between FDs. To visualize these relationships, the distance matrix between all vectors is used as input for principal coordinate analysis (PCoA), which scales data to two dimensions (28). We first examined the position specificity of the FDs by comparing recently-circulating strains from the four clades. Only FDs of positions that evolved from a PNGS in their inferred clade ancestor were compared. Clear clustering of the FDs for the same position was observed (see P values for position-specificity of the patterns in **Fig. 3A** and approach to calculations in the Methods Section). Some positions (e.g., 339) have diversified significantly from the ancestral form (see lines between “Ancestral State” that represents an FD composed only of PNGSs, and the centroid of all clade FDs for each position). Other positions (e.g., 289) have diversified less (i.e., their population-level profiles are composed of a limited number of residues other than the PNGS motif). The FDs of positions 386 and 392, which are directly adjacent on the Env trimer (**Fig. 1A**), were clearly distinct. In addition to position-specificity, we also observed clear clade-specific patterns (**Fig. 3B**). At positions 289, 295, 332 and 339, FDs from the same clade but different regions were clustered (see indication of P values for clade-specificity in each panel of **Fig. 3B**). Clustering patterns were observed at positions 386 and 392, but did not reach statistical significance (consistent with the greater similarity between the clade profiles, **Fig. 2B**). We extended our analyses to compare FDs of the 13 positions in gp120 that contain a PNGS motif in the ancestors of all four clades (**Table S2**). FDs exhibited various degrees of ‘migration’ from the ancestral sequence and position-specific clustering patterns (**Fig. 3C**).

We further analyzed the changes in positional FDs for clades that did not contain a PNGS in their ancestral sequence (**Fig. 4A**). Position 339 in CRF01_AE evolved from an ancestral Asn to an FD similar to the PNGS-derived profiles of position 339 (P value of 0.03 in the permutation test). Of the 17 sites tested, the CRF01_AE FD at position 339 was closest to the centroid of the PNGS-derived FDs of position 339 (i.e., ranked first in proximity; *r=1* in **Fig. 4A** and **Fig. S6A)**. Similar frequencies of residues were observed at this position in CRF01_AE and the other clades (see comparison with the clade A1 in **Fig. 4B**, and all four positions in **Fig. S6B**). The centroid of PNGS-derived FDs for position 332 also ranked first in proximity to the CRF01_AE FD at this position (derived from an ancestral Glu). Greater differences were observed between the PNGS-derived profiles and the FDs at positions 295 in clade C (derived from an ancestral Val, *r=2*) and 289 in clade A1 (derived from an ancestral Thr, *r=6*).

**Fig. 4.**
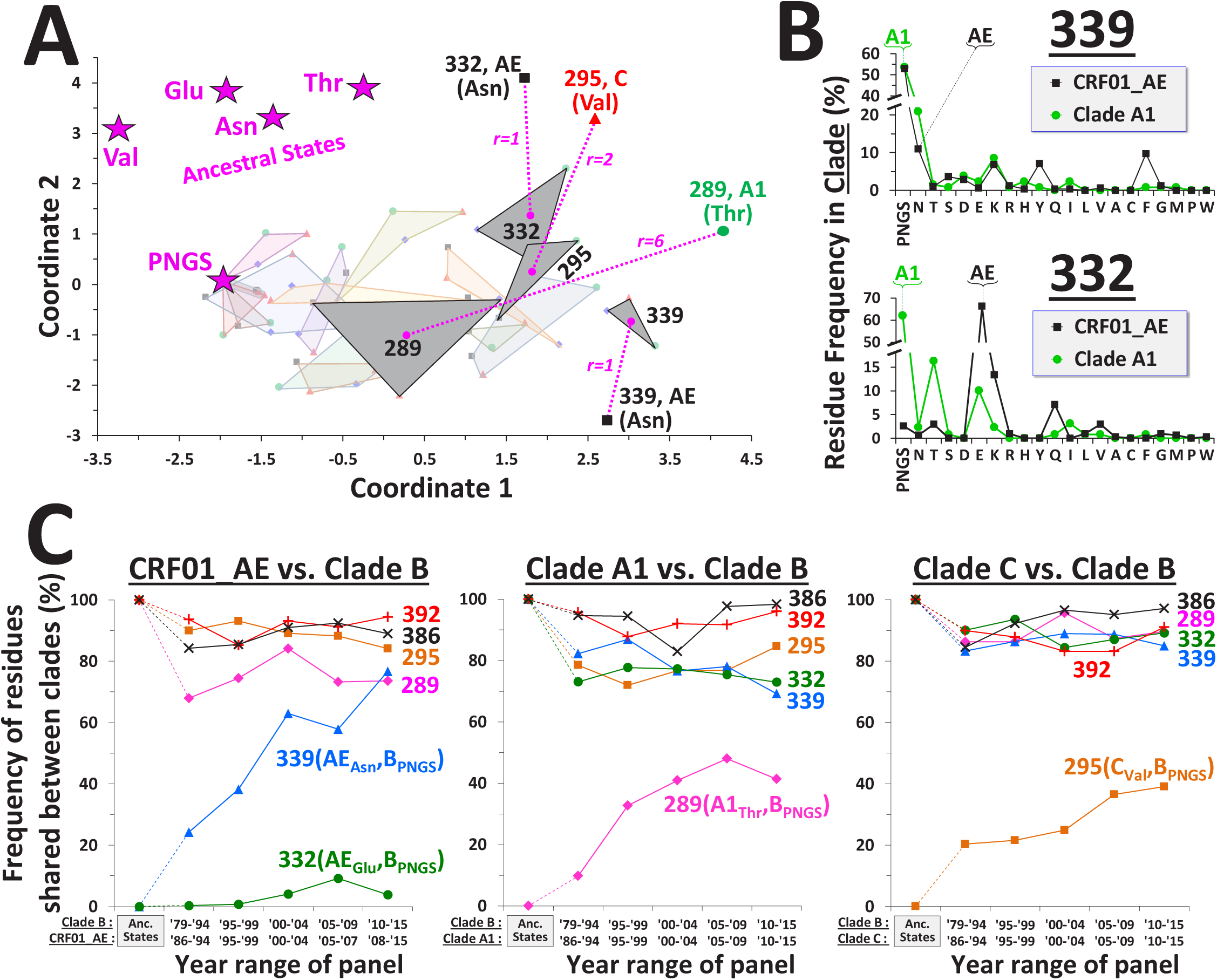
Env positions evolved from distinct clade-ancestral forms toward site-specific FDs, gradually increasing the proportion of shared residues in the diverse clades. **(A)** Relationships between FDs at positions that did not contain an ancestral PNGS motif in one of the four clades. Locations of the clade-ancestral forms (i.e., FDs composed of a single sequence motif) are marked by star symbols. FDs that evolved from a PNGS are connected by grey triangles. FDs that did not evolve from a PNGS are labeled by the lowercase letter of their ancestral residue and connected by dashed lines to the centroid of the PNGS-derived FDs for the same position. Proximity of each FD to the centroid of the corresponding position is indicated by rank order relative to the centroids of all 17 PNGS-derived positions (*r=1* indicates that the corresponding centroid is closest). The thirteen FDs described in **Fig. 3C** are shown in lighter shades. **(B)** FDs at positions 339 and 332 in clade A1 and CRF01_AE. The sequence motifs in the clade ancestors are indicated. **(C)** Historical increase in frequency of residues shared between diverse clades. For any two clades compared, the frequencies of all residues shared between them at each time period were calculated; the sum of the values is shown (see also **Fig. S7**). Residues in the clade ancestors (if both not PNGSs) are indicated. Dotted lines mark changes from the clade-ancestral states.

To understand the impact of the above patterns on sequence similarity at these sites between populations, we examined the historical changes in the proportion of shared residues among the diverse clades. Comparison of the clade B profiles derived from a PNGS with the profiles of other clades that are not derived from a PNGS showed gradual increases in the proportion of shared residues (**Fig. 4C** and **Fig. S7**). The increase in shared PNGSs at these sites accounted for most of the changes (compare with **Fig. 1, B and C**); however, a considerable contribution was also made by other residues that replaced the ancestral forms.

Therefore, distinct Env positions that share the same sequence motif in the clade ancestor have evolved unique FDs of emerging variants. FD profiles are specific for each position (**Figs. 3A** and **3C**) and often for each clade (i.e., conserved in different geographic regions, **Fig. 3B**). Some clades did not contain a PNGS in their ancestral sequence. These clade-founder effects were reduced in most cases by evolution toward the position-specific (PNGS-derived) FD profile. As a consequence of these changes, the proportion of shared residues increased considerably during the course of the pandemic.

### Key positions in the trimer apex evolve toward conserved clade-specific FDs and show gradual reduction of regional founder effects

The V2 variable loop segment at the apex of the Env trimer is targeted by several BNAbs (20–22). This domain has also been linked to vaccine efficacy in the RV144 trial. Presence of anti-V2 antibodies in vaccinees was associated with protection (29). Furthermore, in breakthrough infection analyses of this trial, two signatures of protection were identified, both located in the V2 apex (23). Vaccine efficacy was higher if the infecting strain contained Lys at position 169 or did not contain Ile at 181. We examined residue occupancy at these positions in diverse clades and their evolution during the pandemic. At position 181, the ancestors of clades B and A1 contained Val (associated with protection) whereas the ancestors of clade C and CRF01_AE contained Ile. Interestingly, the frequency of Val in clades B and A1 rapidly decreased during the pandemic and was replaced by Ile (i.e., a form associated with reduced vaccine efficacy, **Fig. 5A**). Similar frequencies of residues were observed in the regional panels of the paraphyletic clade B group (**Fig. 5B** and **Fig. S8A**). Changes in the monophyletic Korean cluster followed the same pattern but lagged behind the paraphyletic group. A distinct but conserved profile was observed for the regional panels of clade C (**Fig. 5B** and **Fig. S8A**). Comparison of the FDs that emerged in clade B at position 181 to other positions that evolved from an ancestral Val revealed the position-specific nature of the profile (**Fig. 5C**).

**Fig. 5.**
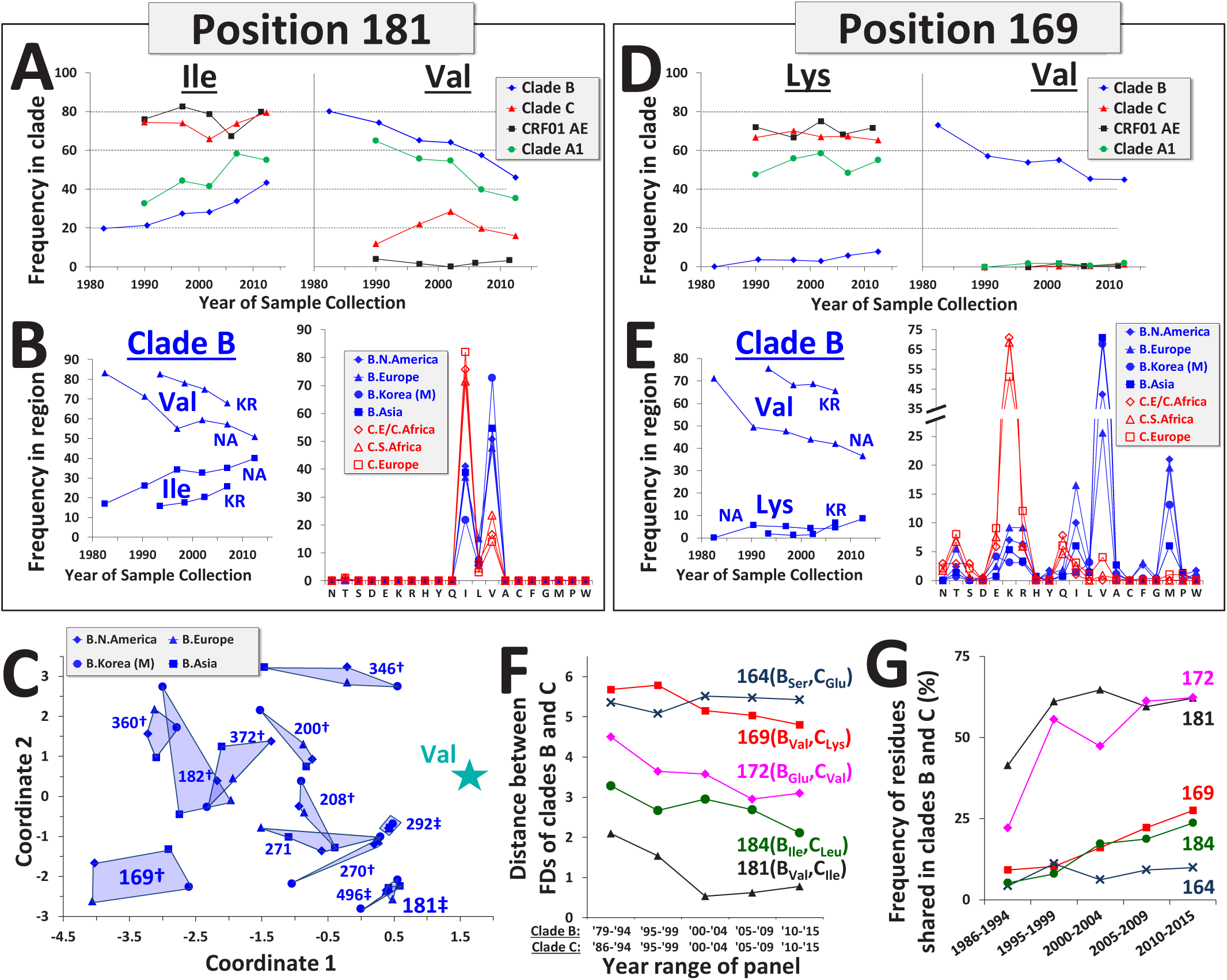
Signature sites of vaccine protection in the RV144 trial have gradually evolved toward conserved clade-specific FDs. **(A,D)** Historical changes in frequency of residues at positions 181 and 169 in diverse clades. **(B,E)** Changes in frequency of residues in clade B viruses from N. America and the monophyletic cluster from Korea (left), and their regional FDs among current strains (right). FDs for all regional panels are shown in **Fig. S8**. **(C)** Relationships between clade B regional FDs at positions that contained a Val residue in the clade-ancestral sequence. Location of the clade-ancestral form (composed only of Val) is indicated by a star symbol. Position-specificity of the patterns is indicated: †, P<0.05; ‡, P<0.005. **(F)** The distance between FDs of clades B and C has gradually declined during the AIDS pandemic. Positions in the V2 loop occupied by distinct residues in the clade B and C ancestors were analyzed. For each time period indicated, the Euclidean distance between the FD vectors in clades B and C was calculated. The clade B period 1979-1986 was compared with the clade C period 1986-1994 due to lack of sufficient clade C sequences from the early 1980’s. Residues in the ancestors of clades B and C are indicated. **(G)** Historical increase in frequency of residues shared between clades B and C for the positions shown in panel F.

At position 169, the inferred ancestors of clades A1, C and CRF01_AE contained Lys (associated with protection) whereas the clade B ancestor contained Val (**Fig. 5D**). The frequency of Val gradually decreased in clade B to less than 50% of circulating strains, and was replaced primarily by Met and Ile, with only a limited increase in Lys (**Fig. 5D**). As expected, the same changes occurred in the monophyletic lineage from Korea and the N. American isolates in the paraphyletic group (**Fig. 5E**). Distinct FD profiles were observed in clades B and C at the most recent time period evaluated, which were conserved in the different regional panels (**Fig. 5E** and **Fig. S8B**).

We asked whether the distance between FDs of distinct clades has changed during the pandemic. Positions in the V2 loop that contained different residues in the ancestors of clades B and C were examined. Sites minimally affected by insertions or deletions were selected. We first calculated for each position the distance between FDs of the paraphyletic groups of these clades at different time periods (**Fig. 5F** and **Fig. S9**). For some positions (e.g., 181 and 172), the inter-clade distance gradually declined. Other positions showed less or no change in this distance (e.g., position 164). Consistent with these results, a gradual increase was observed in the proportion of shared residues in the two clades at each position (see **Fig. 5G**). At some positions, the level of residue similarity appears to have stabilized, whereas at other positions it continues to increase. Therefore, clade-founder effects have gradually decreased during the epidemic, for each position at a different rate.

We also examined whether founder effects are reduced at the regional level when the lineage-founder virus was introduced more recently into a population. Three differences were identified in the V2 segment of the trimer apex between the ancestral sequences of the monophyletic clade B cluster in Korea and the paraphyletic clade B group (see labeled positions in **Fig. 6A** and FDs for all 28 positions in this segment in **Fig. S10**). Significant changes occurred in FDs at the three positions during the pandemic. At position 161, the Korean lineage retained the ancestral Val whereas the frequency of this residue gradually increased in N. America and Europe to replace the ancestral Ile (**Fig. 6B**). Consequently, the FDs at position 161 among currently-circulating strains in the paraphyletic group is evolving toward the FD of the Korean lineage. At position 167, the major changes occurred in the Korean lineage; the ancestral Asn was replaced by Asp and reached a frequency comparable to that in N. America and Europe (**Fig. 6C**). Similar FDs were observed at this position for the different regional groups. An interesting pattern was observed at position 164, which was occupied by Ser in the ancestor of the clade B paraphyletic group but Asn in the founder of the Korean lineage. In all three regions, the frequency of Ser gradually evolved toward a value of 29-34% whereas frequency of Asn evolved toward a value of 20-30% (**Fig. 6D**). Again, similar FDs were observed at position 164 in the different regional groups. Comparison of clade B regional FDs at positions 164 and 167 among recent strains showed clear changes toward the position- and clade-specific FDs (**Fig. 6E**).

**Fig. 6.**
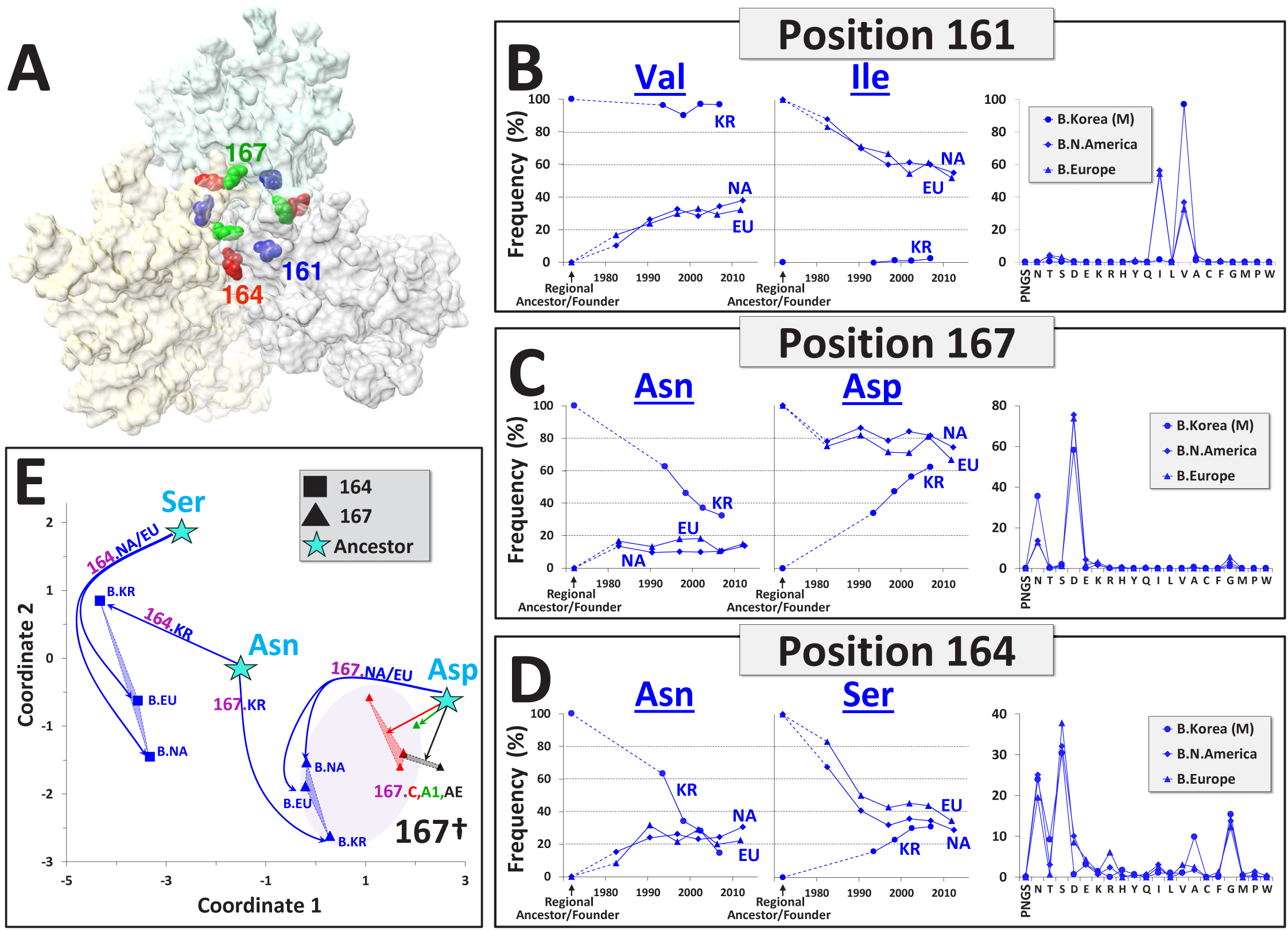
Key positions in the V2 apex have rapidly evolved in clade B from distinct residues in the regional ancestral/founder viruses toward similar FDs. **(A)** Top view of the Env trimer apex (PDB ID 4TVP). Residues in the V2 loop that differ between the clade B regional founder of the monophyletic cluster in Korea and the clade B ancestral sequence are labeled. **(B-D)** Historical changes in frequency of residues at the indicted positions in Korea, N. America and Europe. FDs among current strains in the three regions are shown to the right. Dashed lines indicate the change from the ancestral/founder form of each group. **(E)** Relationships between clade B regional FDs at positions 164 and 167, calculated among recently-circulating viruses. Positions of the clade-ancestral forms (composed of a single residue) are indicated by star symbols. Lines are drawn from the ancestral form to each of the FDs that emerged from that residue. Dotted lines connect between positional FDs of the same position and clade. FDs at position 167 for clades A1, C and CRF01_AE are shown for comparison. Clade-specificity of the pattern was calculated for position 167 as described above: †, P<0.05;.

These results reveal the dominant nature of the forces that guide evolution of each site on Env. They are highly conserved in different populations worldwide and sufficiently strong to decrease clade- and regional-founder effects. Evolution of Env toward well-defined FDs results in a gradual and considerable decrease in the inter-population diversity of this protein.

## Discussion

The Env protein of HIV-1 is tremendously diverse in each population it has infected worldwide (3, 4, 8). The wide range of circulating forms poses a significant hurdle to development of an effective vaccine. Several studies have shown that previously-conserved epitopes on Env are gradually lost, each at a distinct rate (12–14). However, to our knowledge, there has been no report of any clear ‘directionality’ to the changes at a population level (i.e., toward a specific structural form). Here we show that HIV-1 *does* evolve at a population level toward defined ‘target’ states, which are specific for each position of the molecule and often for each lineage of the virus. Selection pressures drive Env toward conserved *distributions of variants*. Key antigenic sites and signatures of vaccine protection have rapidly changed during the pandemic toward well-defined FDs of residues. The pressures are sufficiently strong to overcome founder effects of the virus in different clades and regions. The rapid reduction of clade- and regional-founder effects is surprising but also provides an optimistic outlook on the future of AIDS vaccines.

### The selective pressures that guide evolution of Env are conserved in different geographic regions and virus lineages

The replication machinery of RNA viruses is prone to errors. Random mutation events continuously introduce changes in sequence. Persistence of each variant within the host is determined by constraints applied on RNA secondary structure and by selective pressures applied on the protein, on its fitness and sensitivity to the host immune response. Establishment of the persistent variants in the population is determined by the bottlenecks applied during transmission and conservation of all selective pressures among different hosts (30–33). Our findings suggest that the combined effect of the stochastic events that generate the mutants and the above deterministic forces is a conserved distribution of forms that is specific for each position. Such forces are sufficiently dominant to produce similar profiles in distinct lineages of the virus, even for the lowest-frequency (1-3%) residues. Therefore, in contrast to the in-host environment, which is dominated by stochastic changes (12, 34, 35), the population-level distribution of circulating forms is dominated by conserved deterministic forces.

What are the selective forces that have guided these population-level changes? Fitness pressure is likely the primary form of selection (36–38). As such, frequencies of residues may describe their mean relative fitness in all structural ‘contexts’ within a clade. Clade-specific patterns can thus reflect unique structural properties of their Envs, which present specific fitness constraints (9, 10). Immune pressure applied by antibodies commonly elicited in the infected individual may also cause population-level changes (39–41). For example, it is tempting to hypothesize that rapid replacement of Asn at position 339 of CRF01_AE by a PNGS (**Fig. 1C**) resulted from higher resistance of the latter to neutralization by antibodies (24, 42–44). The observed clade-specific patterns may thus also reflect the unique antigenicity profiles of Envs in each clade (45). Analyses of the relative fitness and neutralization sensitivity of variants at evolving sites of Env will reveal the nature of the pressures that guide population-level changes.

### Design of immunogens against an evolving distribution of forms

HIV-1 has diversified from the group M founder virus to create distinct lineages. Within each, the virus has continued to change in sequence and antigenic properties (9–12). Here we compare evolutionary patterns of Env *between* different populations. Clear changes are observed from distinct ancestral/founder forms in different populations toward similar distributions of residues. For many positions of Env, the virus has evolved from distinct clade-ancestral residues to the current distribution of forms, in which more than 50% of residues are shared. Therefore, diversity has *increased* at the within-population level, whereas it has *decreased* at the between-population level. In some cases, the population-level frequencies of residues appear to have stabilized, such as the PNGS frequencies in the six sites of gp120 (**Fig. 1B**) or the dominant residues at position 164 in the V2 loop apex (**Fig. 6**). In other cases, such as the signatures of vaccine protection, changes appear to progress at historically-constant rates at the clade and regional levels (**Fig. 5**). Current patterns are clearly affected by the time allowed for the changes to occur. For example, the monophyletic clade B lineage in Korea, which dates to the 1960’s (5), was likely introduced into this region in the late 1980’s or early 1990’s. Accordingly, this lineage shows similar patterns of change (**Figs. 3** and **5**) that are delayed relative to other clade B groups (**Fig. 1D** and **Fig. 6**).

Major population-level shifts have occurred in Env antigenicity during the AIDS pandemic (11–14). At several positions located in conserved domains, only a minority of currently-circulating strains contain the clade-ancestral residue. We observe particularly significant changes in clade B at the trimer apex, a domain targeted by multiple quaternary-specific BNAbs including CAP256-VRC26, PG9, PG16 and PGT145 (46–50). Such patterns correspond with the declining breadth of these antibodies; a recent population-level study in the United States showed that nearly half of clade B strains are poorly recognized by PG9 and PG16 (12). Analyses of changes that follow founder events in newly-infected regions allow us to identify the ‘preferred’ forms at each site. Here we focus on the clade B lineage in Korea. At some positions, viruses from the paraphyletic group rapidly gained the Korean founder residue (e.g., position 161, **Fig. 6B**). At other positions, the opposite pattern was observed with rapid reduction of the founder effect (e.g., position 167, **Fig. 6C**). Notably, some positions have evolved toward FDs that are not dominated by any single residue (e.g., position 164 in **Fig. 6D** or 170 and 183 in **Fig. S10**). Conserved patterns of change from each ancestral/founder form allow us to tailor immunogens to the current distribution of residues in each population, the historical time point the infection was established, and the changes expected to occur toward the site-specific profile.

## Methods

### Analyses of HIV-1 Env sequences

All HIV-1 *env* sequences were obtained from the Los Alamos National Lab (LANL) database using the sequence search interface (https://www.hiv.lanl.gov) and from the National Center for Biotechnology Information (NCBI) database (https://www.ncbi.nlm.nih.gov). Non-functional Envs were removed, as were sequences with nucleotide ambiguities or large deletions in conserved regions. A single *env* from each patient and a single sequence from known transmission pairs. In addition, a minimal nucleotide distance of 0.03 nucleotide substitutions per site was applied as a cutoff for selection. **Datasets S1-S4** contain amino acid sequences of all Envs applied in this study. For phylogenetic analyses, nucleotide sequences were aligned using a Hidden Markov Model with the HMMER3 software (51). Phylogenetic trees were reconstructed using the maximum likelihood method using PhyML3 (52). All Env positions described in the manuscript conform to the standard HXBc2 numbering of the Env protein (53). Potential N-linked glycosylation sites (PNGSs) were defined by the presence of the sequence Asn-X-Ser/Thr, where X can be any amino acid except Pro.

### Statistical analyses of residue frequency distributions and specificity of the patterns for position and clade

Frequency distributions (FDs) describe the percent occupancy of Env positions by each residue in a defined population. Each FD is a vector composed of 20 or 21 features (20 amino acids and a PNGS). Residues with frequencies lower than 0.75% (for regional panels) or lower than 0.6% (for whole-clade panels) were assigned a value of 0.1, and values were log_10_-transformed. Log_10_ conversion of frequency values increases the relative contribution of minor variants to the profile. To calculate relationships between FDs, Euclidean distances between all 21-feature vectors were measured. For graphical representation of the relationships between FDs, the distance matrix between their vectors was used as input for the Torgerson scaling method (28).

To determine the clade-specificity of FDs, we first calculated for each position (*p*) the coordinates of the centroid (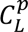) among vectors from the same clade (*L*). For each clade, the mean intra-clade distance (*d_intra–clade_*) was calculated as the average Euclidean distance between 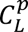 and all regional vectors of the same clade (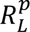), formally 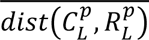. In addition, we calculated mean inter-clade distance (*d_inter–clade_*) as the average Euclidean distance between the centroid of clade *L* and all other clade centroids 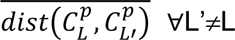. We define the 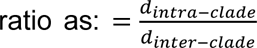. The baseline ratio (S_base_) was calculated as the *ratio* using all panels. Then, within the position whose clade specificity is being calculated, clade identifiers were permuted and randomly assigned to each panel, from which the permuted ratio (S_rand_) was calculated. The permutation process was repeated 10,000 times. The P value was calculated. The permutation process was repeated 10,000 times. The P value was calculated as 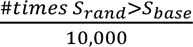. To establish the position specificity of the profiles, the centroid of all regional profiles for a given position (p) was calculated (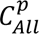). Here intra-position distance (*d_intra–position_*) was calculated as the average Euclidean distance between 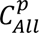 and all profiles of position *p* (for all clades and regions), more formally 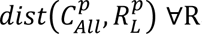, L where 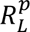 denotes the profile in region R with position p and clade L. Then, inter-position distance (*d_inter–position_*) was calculated as the average Euclidean distance between 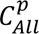 and all other positional centroids, more formally 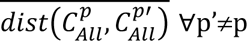. Finally, the ratio was determined as: 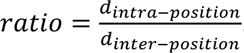. Similar to clade specificity calculations, baseline ratio (S_base_) was first calculated. Position identifiers were then shuffled and randomly assigned to each panel, from which the permuted ratio (S_rand_) was calculated. The shuffling process was repeated 10,000 times, and the P value was calculated as 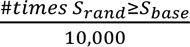.

Source code for calculating residue FDs, position specificity and clade specificity (including all datasets applied in this work) can be found at https://github.com/haimlab/HIV.

## Acknowledgments

We thank Kevin Rice, Ernesto Fuentes, Lokesh Gakhar and Andrés Finzi for helpful discussions, and Wendy Maury and Stanley Perlman for critical reading of this manuscript.

## References

1. Coffin JM (1995) HIV population dynamics in vivo: implications for genetic variation, pathogenesis, and therapy. Science 267(5197):483–489.

2. Preston BD, Poiesz BJ, & Loeb LA (1988) Fidelity of HIV-1 reverse transcriptase. Science 242(4882):1168–1171.

3. Buonaguro L, Tornesello ML, & Buonaguro FM (2007) Human immunodeficiency virus type 1 subtype distribution in the worldwide epidemic: pathogenetic and therapeutic implications. J Virol 81(19):10209–10219.

4. Taylor BS, Sobieszczyk ME, McCutchan FE, & Hammer SM (2008) The challenge of HIV-1 subtype diversity. N Engl J Med 358(15):1590–1602.

5. Kim GJ, et al. (2012) Estimating the origin and evolution characteristics for Korean HIV type 1 subtype B using Bayesian phylogenetic analysis. AIDS Res Hum Retroviruses 28(8):880–884.

6. Shen C, Craigo J, Ding M, Chen Y, & Gupta P (2011) Origin and dynamics of HIV-1 subtype C infection in India. PLoS One 6(10):e25956.

7. Neogi U, et al. (2012) Molecular epidemiology of HIV-1 subtypes in India: origin and evolutionary history of the predominant subtype C. PLoS One 7(6):e39819.

8. Korber B, et al. (2001) Evolutionary and immunological implications of contemporary HIV-1 variation. Br Med Bull 58:19–42.

9. Lynch RM, Shen T, Gnanakaran S, & Derdeyn CA (2009) Appreciating HIV type 1 diversity: subtype differences in Env. AIDS Res Hum Retroviruses 25(3):237–248.

10. Gnanakaran S, et al. (2007) Clade-specific differences between human immunodeficiency virus type 1 clades B and C: diversity and correlations in C3-V4 regions of gp120. J Virol 81(9):4886–4891.

11. Hraber P, et al. (2014) Impact of clade, geography, and age of the epidemic on HIV-1 neutralization by antibodies. Journal of virology 88(21):12623–12643.

12. DeLeon O, et al. (2017) Accurate predictions of population-level changes in sequence and structural properties of HIV-1 Env using a volatility-controlled diffusion model. PLoS Biol 15(4):e2001549.

13. Bouvin-Pley M, et al. (2013) Evidence for a continuous drift of the HIV-1 species towards higher resistance to neutralizing antibodies over the course of the epidemic. PLoS pathogens 9(7):e1003477.

14. Bunnik EM, et al. (2010) Adaptation of HIV-1 envelope gp120 to humoral immunity at a population level. Nature medicine 16(9):995–997.

15. Gaschen B, et al. (2002) Diversity considerations in HIV-1 vaccine selection. Science 296(5577):2354–2360.

16. Korber B, Hraber P, Wagh K, & Hahn BH (2017) Polyvalent vaccine approaches to combat HIV-1 diversity. Immunol Rev 275(1):230–244.

17. Korber B & Gnanakaran S (2009) The implications of patterns in HIV diversity for neutralizing antibody induction and susceptibility. Curr Opin HIV AIDS 4(5):408–417.

18. Cao L, et al. (2017) Global site-specific N-glycosylation analysis of HIV envelope glycoprotein. Nat Commun 8:14954.

19. Lyumkis D, et al. (2013) Cryo-EM structure of a fully glycosylated soluble cleaved HIV-1 envelope trimer. Science 342(6165):1484–1490.

20. Lee JH, et al. (2017) A Broadly Neutralizing Antibody Targets the Dynamic HIV Envelope Trimer Apex via a Long, Rigidified, and Anionic beta-Hairpin Structure. Immunity 46(4):690–702.

21. Walker LM, et al. (2009) Broad and potent neutralizing antibodies from an African donor reveal a new HIV-1 vaccine target. Science 326(5950):285–289.

22. Doria-Rose NA, et al. (2016) New Member of the V1V2-Directed CAP256-VRC26 Lineage That Shows Increased Breadth and Exceptional Potency. J Virol 90(1):76–91.

23. Rolland M, et al. (2012) Increased HIV-1 vaccine efficacy against viruses with genetic signatures in Env V2. Nature 490(7420):417–420.

24. Kong L, et al. (2013) Supersite of immune vulnerability on the glycosylated face of HIV-1 envelope glycoprotein gp120. Nature structural & molecular biology 20(7):796–803.

25. Krumm SA, et al. (2016) Mechanisms of escape from the PGT128 family of anti-HIV broadly neutralizing antibodies. Retrovirology 13:8.

26. Pritchard LK, et al. (2015) Glycan clustering stabilizes the mannose patch of HIV-1 and preserves vulnerability to broadly neutralizing antibodies. Nat Commun 6:7479.

27. Travers SA (2012) Conservation, Compensation, and Evolution of N-Linked Glycans in the HIV-1 Group M Subtypes and Circulating Recombinant Forms. ISRN AIDS 2012:823605.

28. Torgerson WS (1958) Theory and methods of scaling (Wiley, New York,) p 460 p.

29. Haynes BF, et al. (2012) Immune-correlates analysis of an HIV-1 vaccine efficacy trial. The New England journal of medicine 366(14):1275–1286.

30. Claiborne DT, et al. (2015) Replicative fitness of transmitted HIV-1 drives acute immune activation, proviral load in memory CD4+ T cells, and disease progression. Proceedings of the National Academy of Sciences of the United States of America 112(12):E1480–1489.

31. Keele BF, et al. (2008) Identification and characterization of transmitted and early founder virus envelopes in primary HIV-1 infection. Proceedings of the National Academy of Sciences of the United States of America 105(21):7552–7557.

32. Rademeyer C, et al. (2016) Features of Recently Transmitted HIV-1 Clade C Viruses that Impact Antibody Recognition: Implications for Active and Passive Immunization. PLoS pathogens 12(7):e1005742.

33. Salazar-Gonzalez JF, et al. (2009) Genetic identity, biological phenotype, and evolutionary pathways of transmitted/founder viruses in acute and early HIV-1 infection. J Exp Med 206(6):1273–1289.

34. Kouyos RD, Althaus CL, & Bonhoeffer S (2006) Stochastic or deterministic: what is the effective population size of HIV-1? Trends in microbiology 14(12):507–511.

35. Merrill SJ (2005) The stochastic dance of early HIV infection. J Comput Appl Math 184(1):242–257.

36. Haddox HK, Dingens AS, Hilton SK, Overbaugh J, & Bloom JD (2018) Mapping mutational effects along the evolutionary landscape of HIV envelope. Elife 7.

37. Zanini F, Puller V, Brodin J, Albert J, & Neher RA (2017) In vivo mutation rates and the landscape of fitness costs of HIV-1. Virus Evol 3(1):vex003.

38. Bons E, Bertels F, & Regoes RR (2018) Estimating the mutational fitness effects distribution during early HIV infection. Virus Evol 4(2):vey029.

39. Gray ES, et al. (2009) Antibody specificities associated with neutralization breadth in plasma from human immunodeficiency virus type 1 subtype C-infected blood donors. Journal of virology 83(17):8925–8937.

40. Georgiev IS, et al. (2013) Delineating antibody recognition in polyclonal sera from patterns of HIV-1 isolate neutralization. Science 340(6133):751–756.

41. Walker LM, et al. (2010) A limited number of antibody specificities mediate broad and potent serum neutralization in selected HIV-1 infected individuals. PLoS pathogens 6(8):e1001028.

42. Pejchal R, et al. (2011) A potent and broad neutralizing antibody recognizes and penetrates the HIV glycan shield. Science 334(6059):1097–1103.

43. Reitter JN, Means RE, & Desrosiers RC (1998) A role for carbohydrates in immune evasion in AIDS. Nat Med 4(6):679–684.

44. Wei X, et al. (2003) Antibody neutralization and escape by HIV-1. Nature 422(6929):307–312.

45. Bures R, et al. (2002) Regional clustering of shared neutralization determinants on primary isolates of clade C human immunodeficiency virus type 1 from South Africa. Journal of virology 76(5):2233–2244.

46. Chuang GY, et al. (2013) Residue-level prediction of HIV-1 antibody epitopes based on neutralization of diverse viral strains. J Virol 87(18):10047–10058.

47. McLellan JS, et al. (2011) Structure of HIV-1 gp120 V1/V2 domain with broadly neutralizing antibody PG9. Nature 480(7377):336–343.

48. Doria-Rose NA, et al. (2012) A short segment of the HIV-1 gp120 V1/V2 region is a major determinant of resistance to V1/V2 neutralizing antibodies. J Virol 86(15):8319–8323.

49. Bricault CA, et al. (2019) HIV-1 Neutralizing Antibody Signatures and Application to Epitope-Targeted Vaccine Design. Cell Host Microbe 25(1):59–72 e58.

50. Doria-Rose NA, et al. (2014) Developmental pathway for potent V1V2-directed HIV-neutralizing antibodies. Nature 509(7498):55–62.

51. Gaschen B, Kuiken C, Korber B, & Foley B (2001) Retrieval and on-the-fly alignment of sequence fragments from the HIV database. Bioinformatics 17(5):415–418.

52. Nickle DC, et al. (2007) HIV-specific probabilistic models of protein evolution. PloS one 2(6):e503.

53. Korber B, Foley B, Kuiken C, Pillai S, & Sodroski J (1998) Numbering positions in HIV relative to HXBc2. Los Alamos: Los Alamos Natl. Lab.:iii-102–iii-103.

