## Supporting Information for "Gradual decrease in the inter-population diversity of HIV-1 Env among group M lineages worldwide"

#### Supporting Figures

**Fig. S1.** Phylogenetic trees of Env panels used in this study. Each Env is derived from a distinct patient. **(A-C)** Trees for clade B (1,942 patients), clade C (1,248 patients) and clade A1 (335 patients) were constructed by the maximum likelihood method using nucleotide sequences of the Env segment that spans the region from amino acid position 131 to 511 (for clades B and C), or 156 to 459 (for clade A1). All trees are rooted to the clade ancestor. Amino acid alignments of the gp120 sequences are found in **Datasets S1-S3**. **(D)** The tree for CRF01_AE (543 patients) was constructed using nucleotide sequences of the entire *env* gene. Amino acid alignment of the gp160 sequences is found in **Dataset S4**.

**Fig. S2.** Frequencies of PNGSs at six positions of gp120, calculated among strains recently circulating in the indicated regions (see year ranges for each group in **Table S1**). Regional panels that contained a PNGS motif at the indicated positions in their inferred clade ancestors are shown. The clade B panel from Asia does not include samples from Korea. M, the monophyletic groups of clade B and C viruses from Korea and India/Nepal, respectively. Within-clade comparisons of the positions are shown in **Fig. 1D**. Horizontal bars describe clade averages (among all regional panels) and are colored by clade. Error bars, standard error of the mean (SEM).

**Fig. S3.** Amino acid frequency distribution (FD) profiles at Env positions occupied by a PNGS motif in the ancestral sequences of clades B, C, A1 and CRF01_AE. **(A)** FD profiles for the indicated positions were calculated among viruses of each clade currently circulating worldwide. "N" indicates presence of Asn that is not part of a PNGS motif. **(B,C)** FD profiles in regional panels of currently-circulating strains from clades B and C. **(D)** Positional FDs for clade C Envs from S. Africa and the monophyletic cluster from India and Nepal. The monophyletic lineage of isolates from India and Nepal is relatively small; 61 unique-patient sequences are available for the year range 2000-2015 (relative to 239 in the S. African panel from 2007 to 2015). Nevertheless, the FD profile at many positions is similar to other clade C regional panels. **(E)** Nucleotide sequences in the inferred ancestors of the indicated clades at the first and third residues of the N-X-S/T triplet. Sites with identical nucleotide sequences at the two positions are shaded similarly. Asterisks indicate absence of the sequence motif for a PNGS. Differential synonymous codon usage did not correspond with the patterns of emerging residues. For example, the inferred clade B ancestor contains the same sequence (AAT/ACT) at positions 339/341 and 392/394. Nevertheless, the FDs that evolved at positions 339 and 392 are distinct (**Fig. 2**, **A-C**).

**Fig. S4.** Historical changes in frequencies of residues that replaced the clade-ancestral PNGS motif at six positions of gp120. Values represent the percent frequency of each residue (among all variants) at the indicated positions during the 5-7-year periods. Only positional profiles of clades that contained a PNGS motif in their inferred ancestors are shown. High-frequency residues are labeled by their three-letter codes. "N" and “Asn” indicate an asparagine residue that is not part of a PNGS motif.

**Fig. S5.** Frequencies of residues at six positions in the glycan shield of gp120 in regional panels of Envs from clades B, C, A1 and CRF01_AE. A one-way ANOVA test that compares residue frequencies between positions was performed for clades B and C; P values are color coded as indicated on the right.

**Fig. S6.** Env positions evolved from distinct clade-ancestral forms toward site-specific FD profiles. **(A)** Position-specificity of FDs. Euclidean distances between the four FDs not derived from an ancestral PNGS motif (positions 332, 339, 295 and 289) and the centroids of all 17 PNGS-derived FDs were calculated. Distances are color coded by value. The closest position to each non-PNGS-derived FD is highlighted by a red border. **(B)** FDs at Env positions for which one of the four clades does not contain a PNGS in its ancestral sequence. Values describe frequency of residues among strains that currently circulate worldwide. The sequence motif in each clade ancestor is indicated.

**Fig. S7.** Increase in sequence similarity between clade B and clades C, A1 and CRF01_AE at six positions in the glycan shield of Env. The frequency of residues at the indicated positions was calculated in each clade for different time periods of the pandemic. For any two clades compared, the shared frequency of each residue was determined and the sum of all values calculated. Data are shown for the first and last periods evaluated for each clade. Line graphs to the right show the frequency of residues shared at all time periods.

**Fig. S8.** FDs of residues at positions 181 **(A)** and 169 **(B)**, calculated for regional panels of recently-circulating viruses from the indicated clades. Frequency values for Korea and India/Nepal were calculated among isolates of the monophyletic clusters that circulate in these regions. The clade B panel from Asia does not include isolates from Korea. The residue found in the clade-ancestral sequence is labeled ‘Anc’.

**Fig. S9.** Historical changes in frequency of residues at position 169 in clades B and C. **(A)** Changes in frequency of each residue among all isolates from each clade, calculated for consecutive 5-7-year periods. **(B)** FDs of residues at position 169 in clades B and C, calculated among viruses isolated from samples collected at the indicated time periods.

**Fig. S10.** FDs of residues at positions 156-185 in the V2 loop of Env among clade B isolates that currently circulate in the indicated geographic regions. Frequency values for Korea were calculated for isolates from the monophyletic clade B cluster that circulates in this region.

#### Supporting Datasets

**Dataset S1.** Amino acid alignment (in FASTA format) of clade B gp120 sequences used in this study (1,942 Envs). All Envs are aligned against the HXBc2 isolate. A phylogenetic tree of the sequences is found in **Fig. S1A**.

**Dataset S2.** Amino acid alignment (in FASTA format) of clade C gp120 sequences used in this study (1,248 Envs). Sequences are aligned against the HXBc2 isolate. A phylogenetic tree of the sequences is found in **Fig. S1B**.

**Dataset S3.** Amino acid alignment (in FASTA format) of clade A1 gp120 sequences used in this study (335 Envs). Sequences are aligned against the HXBc2 isolate. A phylogenetic tree of the sequences is found in **Fig. S1C.**

**Dataset S4.** Amino acid alignment (in FASTA format) of CRF01_AE gp160 sequences used in this study (543 Envs). Sequences are aligned against the HXBc2 isolate. A phylogenetic tree of the sequences is found in **Fig. S1D.**

**Fig. S1A**

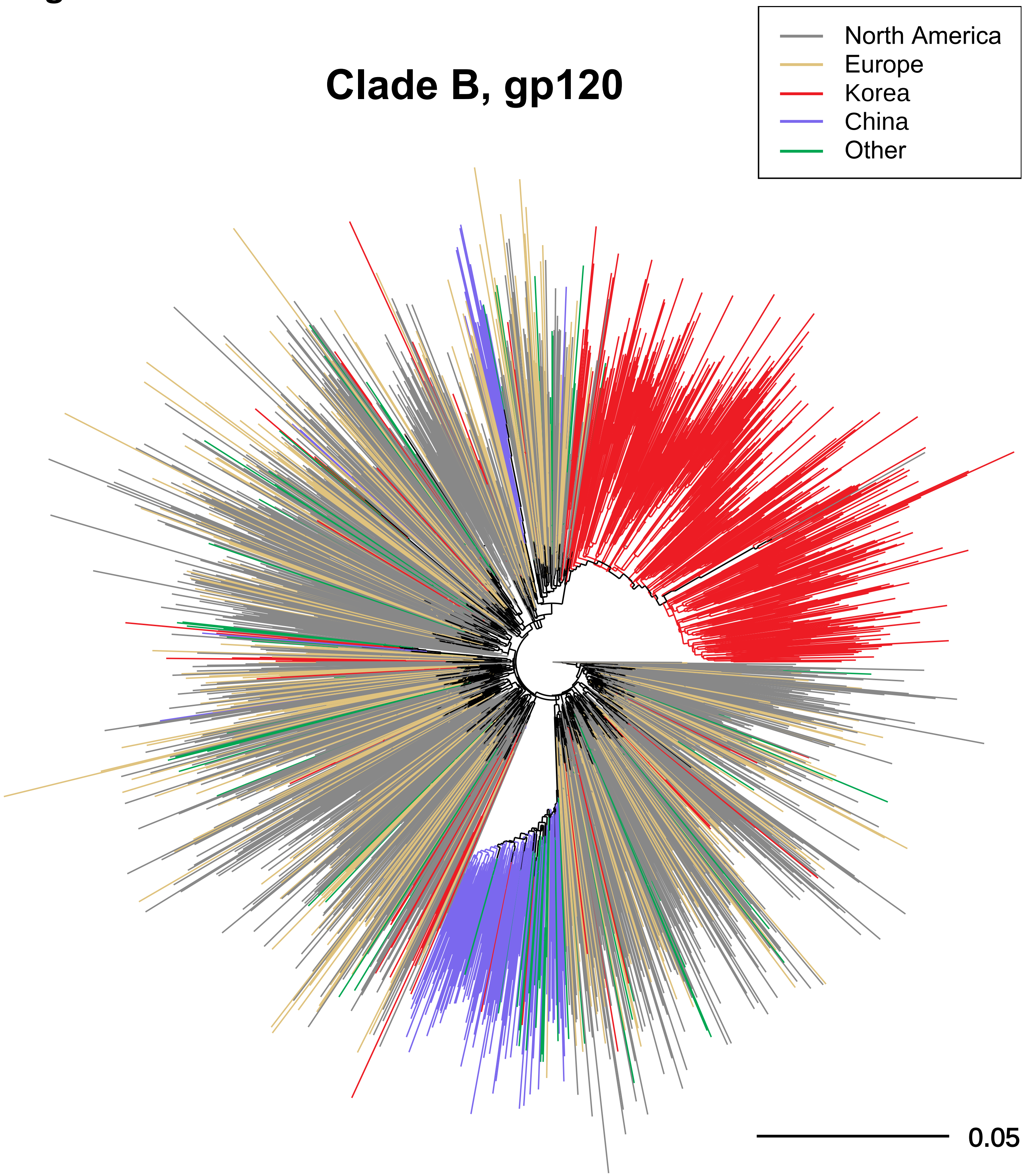

**Fig. S1B**

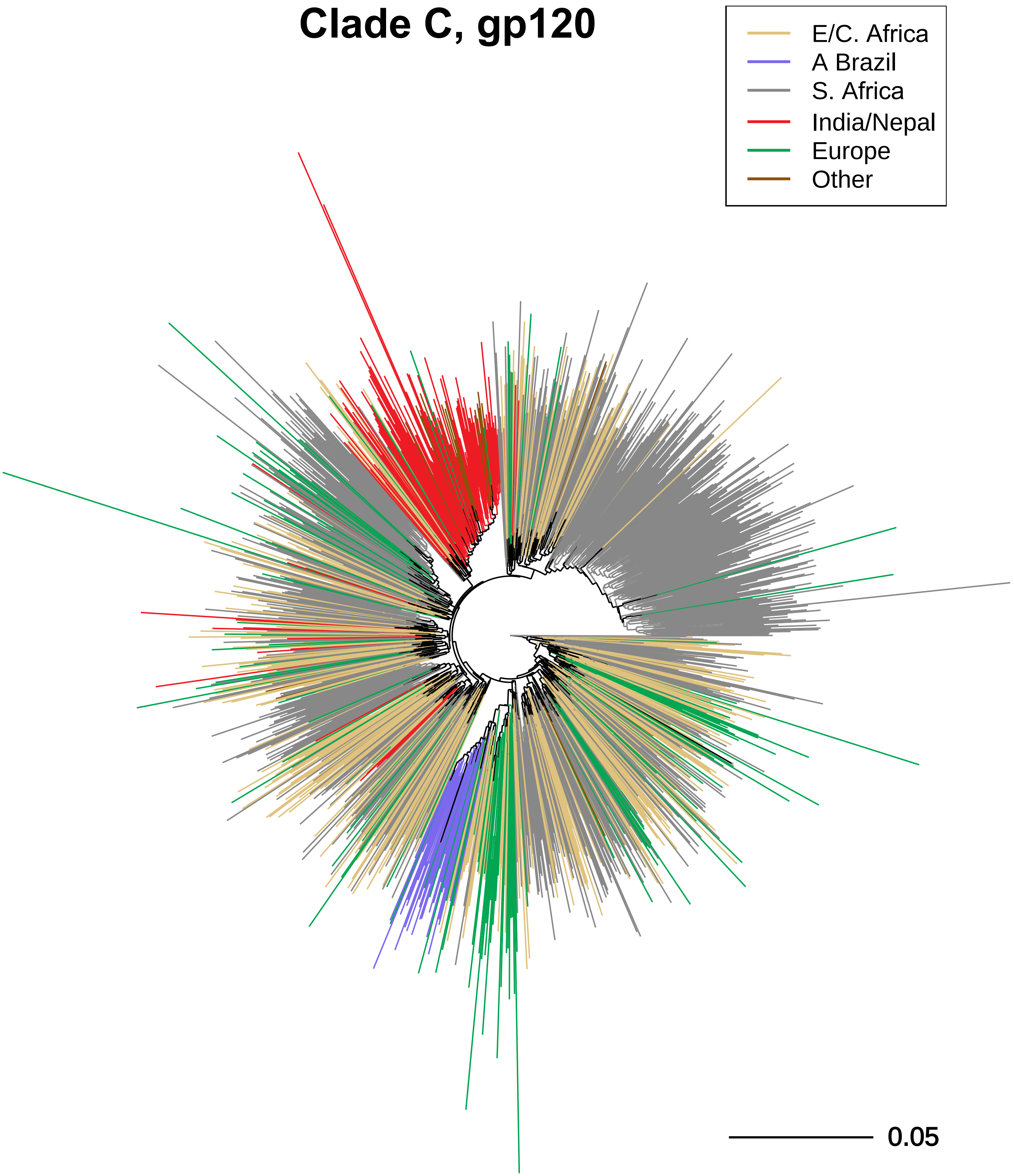
Clade C, gp120
E/C. Africa
A Brazil
S. Africa
India/Nepal
Europe
Other
0.05

**Fig. S1C**

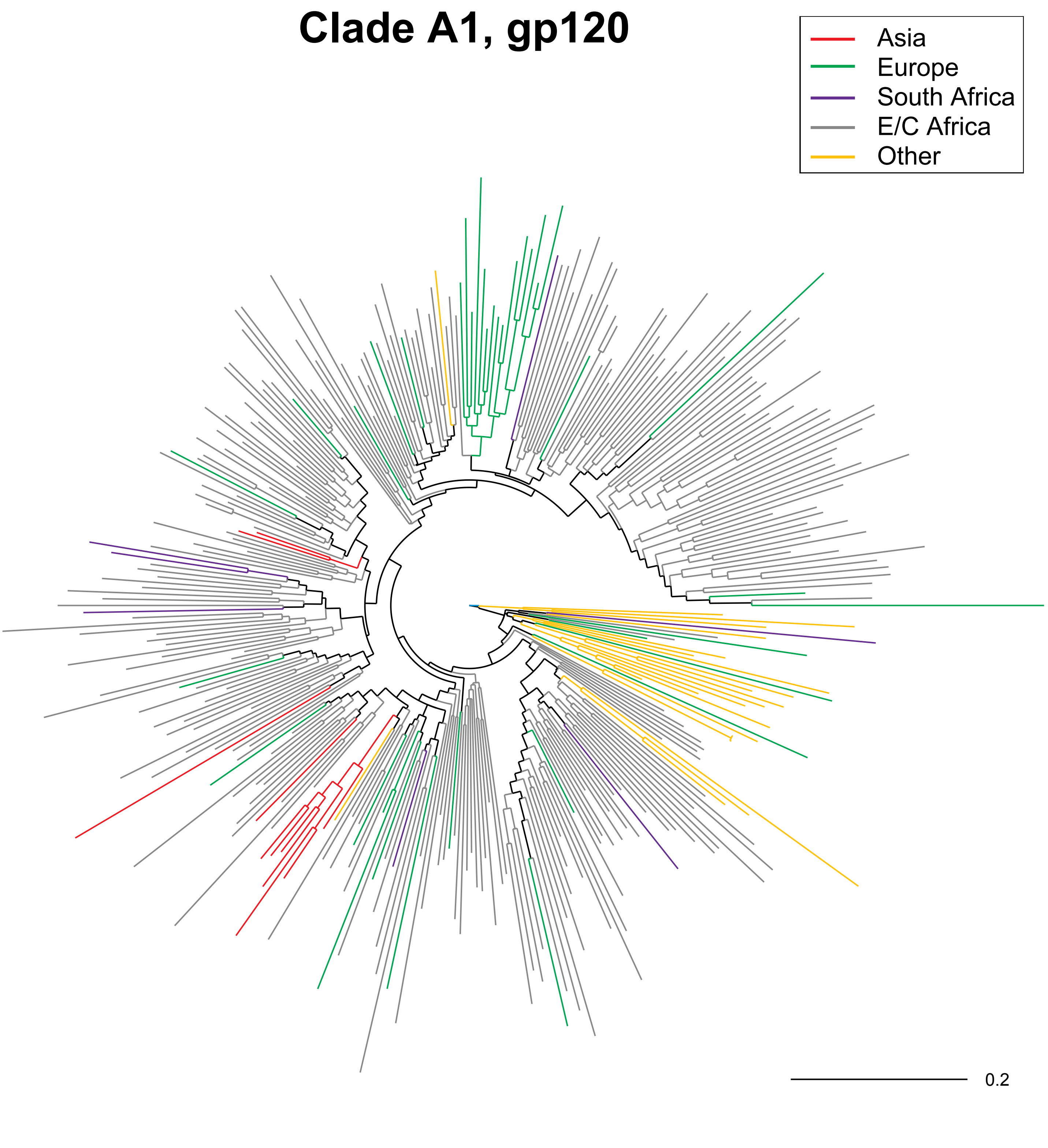

**Fig. S1D**

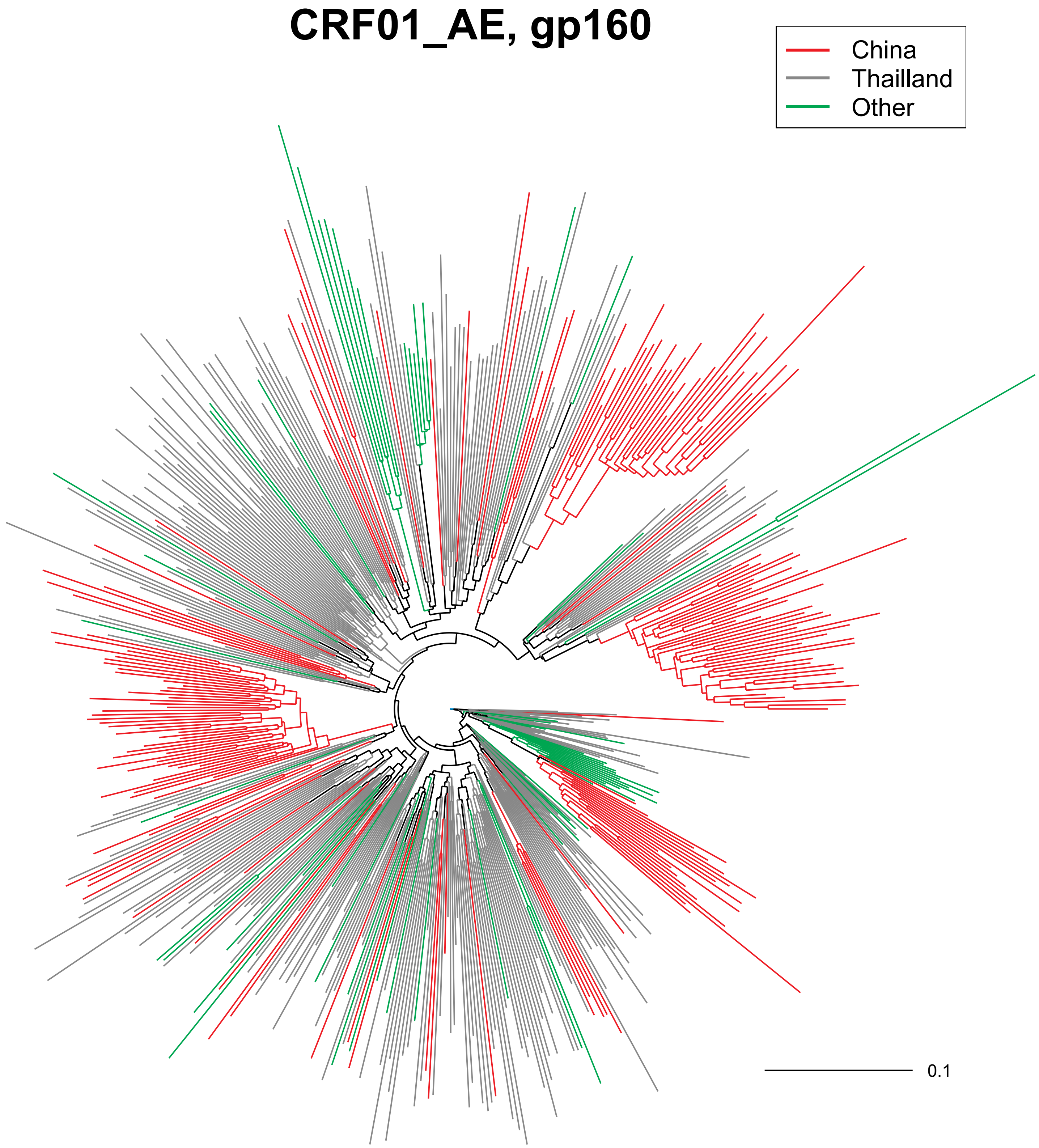

#### Fig. S2

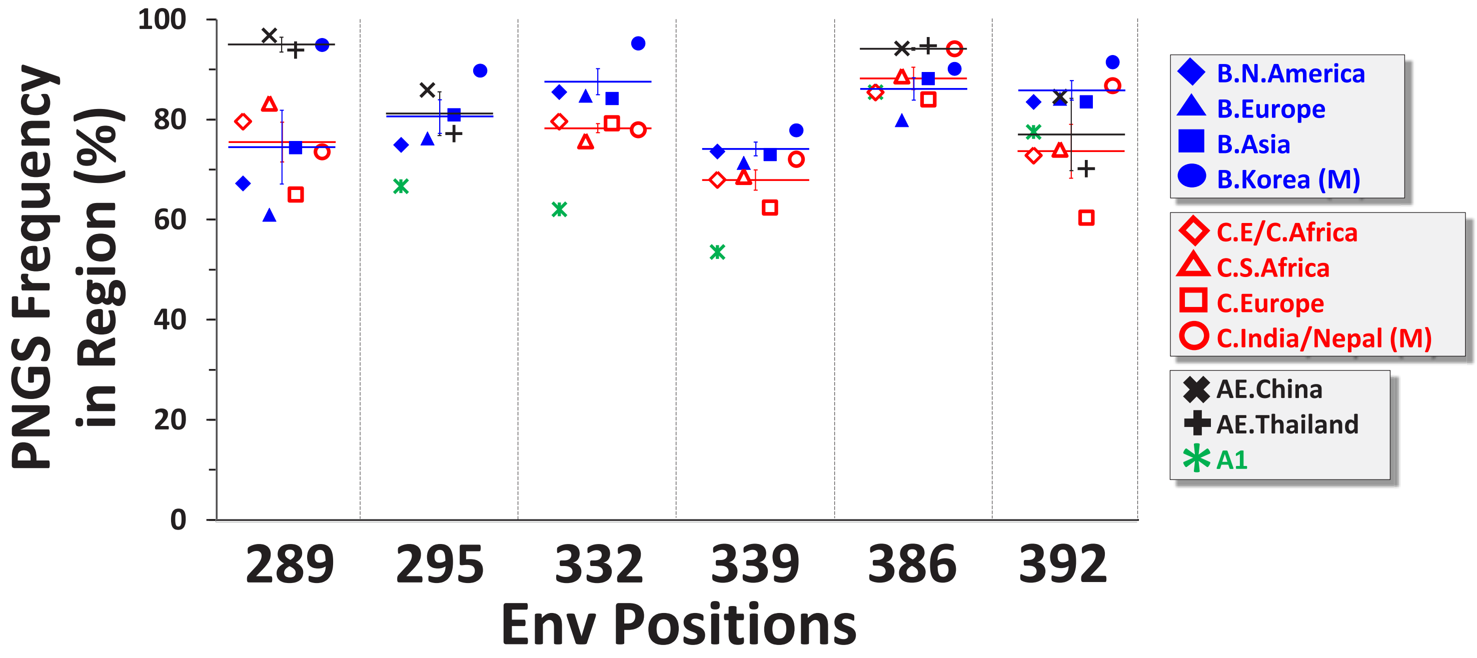
PNGS Frequency in Region (%)
100
80
60
40
20
0
289
295
332
339
386
392
Env Positions
B.N.America
B.Europe
B.Asia
B.Korea (M)
C.E/C.Africa
C.S.Africa
C.Europe
C.India/Nepal (M)
AE.China
AE.Thailand
A1

### Fig. S3

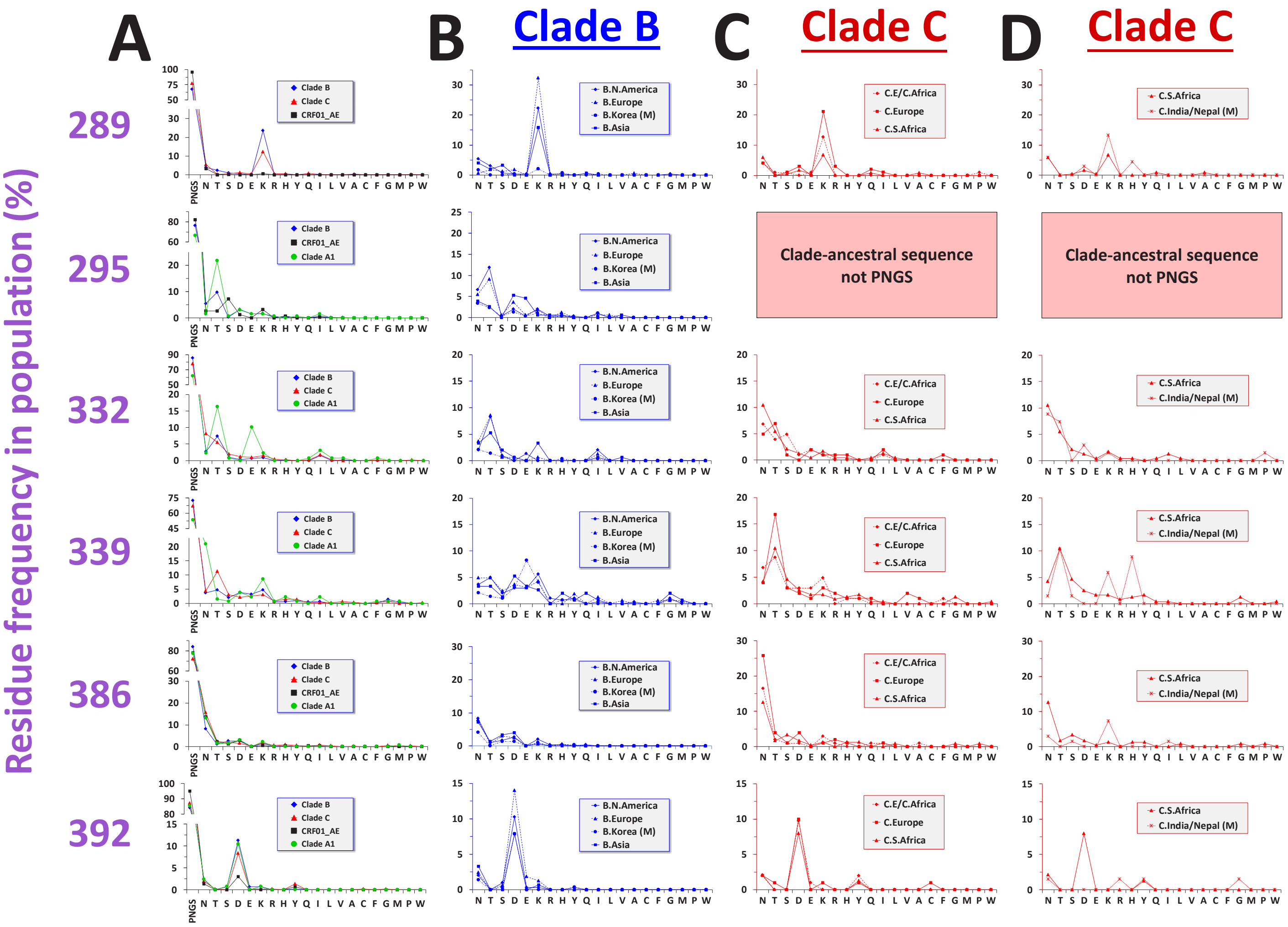

E

| Env Positions | Clades B | A1 | C | AE |
|---|---|---|---|---|
| 289/291 | AAT/TCT | ACC/CCT* | AAT/TCT | AAT/TCT |
| 295/297 | AAT/ACA | AAT/ACC | GTG/ACA* | AAT/ACC |
| 332/334 | AAC/AGT | AAT/AGT | AAC/AGT | GAG/AAT* |
| 339/341 | AAT/ACT | AAT/ACT | AAT/ACT | AAT/GTT* |
| 386/388 | AAT/ACA | AAT/TCA | AAT/TCA | AAT/ACA |
| 392/394 | AAT/ACT | AAT/ACT | AAT/ACA | AAT/ACT |

#### Fig. S4

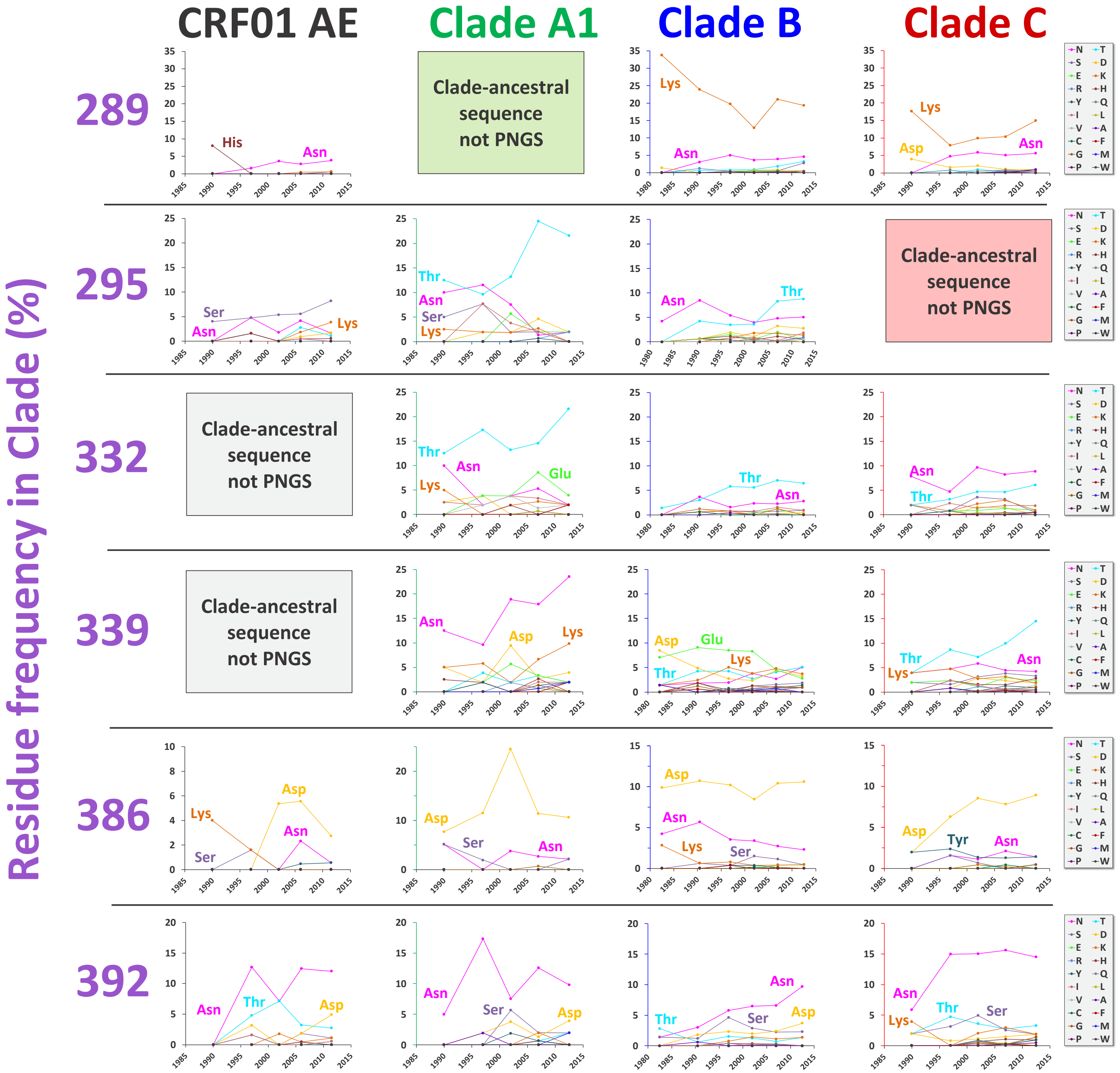
CRF01 AE
Clade A1
Clade B
Clade C
Residue frequency in Clade (%)
289
295
332
339
386
392
Clade-ancestral sequence not PNGS
His
Asn
Lys
Asp
Ser
Thr
Glu
Tyr
N
S
E
R
Y
I
V
C
G
P
T
D
K
H
Q
L
A
F
M
W

### Fig. S5

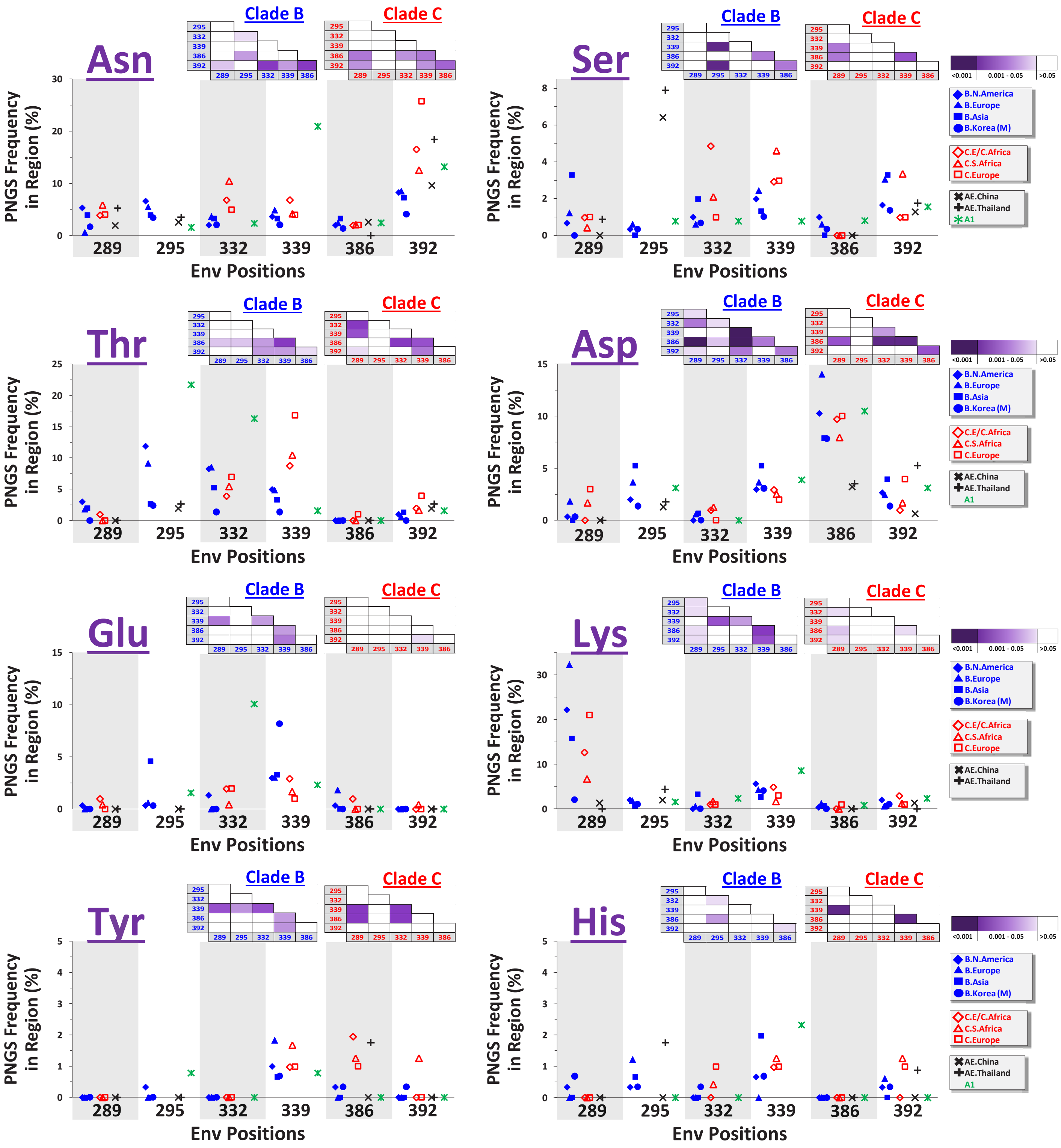
Asn
Ser
Thr
Asp
Glu
Lys
Tyr
His
Clade B
Clade C
PNGS Frequency in Region (%)
Env Positions
289
295
332
339
386
392
<0.001
0.001 - 0.05
>0.05
B.N.America
B.Europe
B.Asia
B.Korea (M)
C.E/C.Africa
C.S.Africa
C.Europe
AE.China
AE.Thailand
A1

**Fig. S6**

# A

| | 88 | 156 | 160 | 197 | 234 | 241 | 262 | 276 | 289 | 295 | 301 | 332 | 339 | 356 | 386 | 392 | 448 |
|---|---|---|---|---|---|---|---|---|---|---|---|---|---|---|---|---|---|
| 339(Asn), CRF01_AE | 4.90 | 4.69 | 3.36 | 4.73 | 3.45 | 4.46 | 4.90 | 4.09 | 3.49 | 3.60 | 4.07 | 4.04 | 2.99 | 3.53 | 4.33 | 3.41 | 3.99 |
| 332(Glu), CRF01_AE | 5.24 | 5.32 | 5.18 | 5.31 | 5.07 | 5.01 | 5.24 | 5.32 | 4.68 | 4.62 | 4.74 | 4.24 | 4.53 | 5.03 | 5.52 | 5.08 | 5.09 |
| 295(Val), Clade C | 5.03 | 4.95 | 4.91 | 5.11 | 4.50 | 4.50 | 5.03 | 4.91 | 4.47 | 3.92 | 4.50 | 3.50 | 4.25 | 4.54 | 5.28 | 4.61 | 4.63 |
| 289(Thr), Clade A1 | 5.46 | 5.34 | 4.39 | 5.21 | 3.94 | 5.01 | 5.46 | 4.80 | 4.18 | 3.76 | 4.68 | 4.39 | 3.85 | 3.59 | 4.79 | 3.93 | 4.30 |

# B

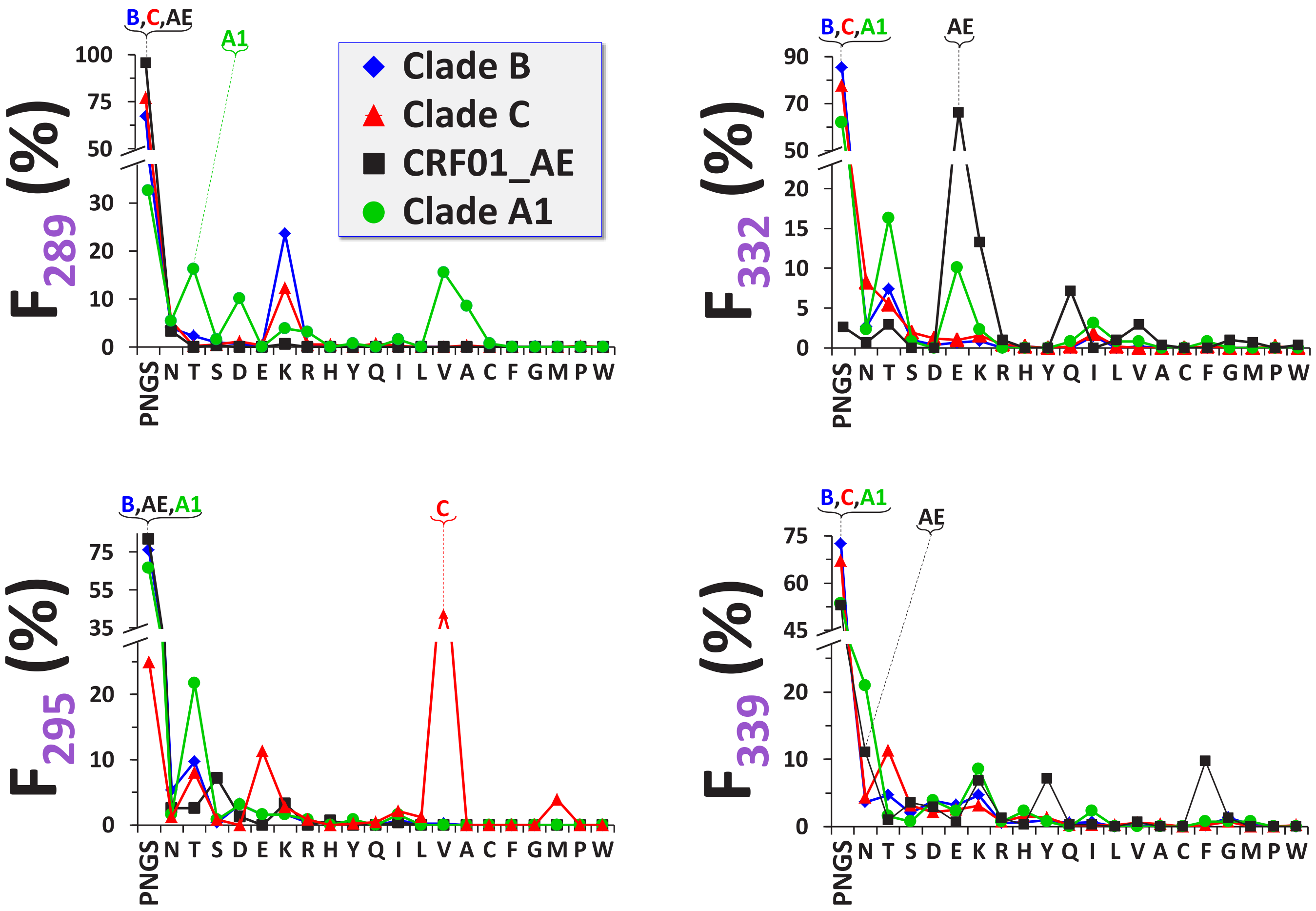

#### Fig. S7

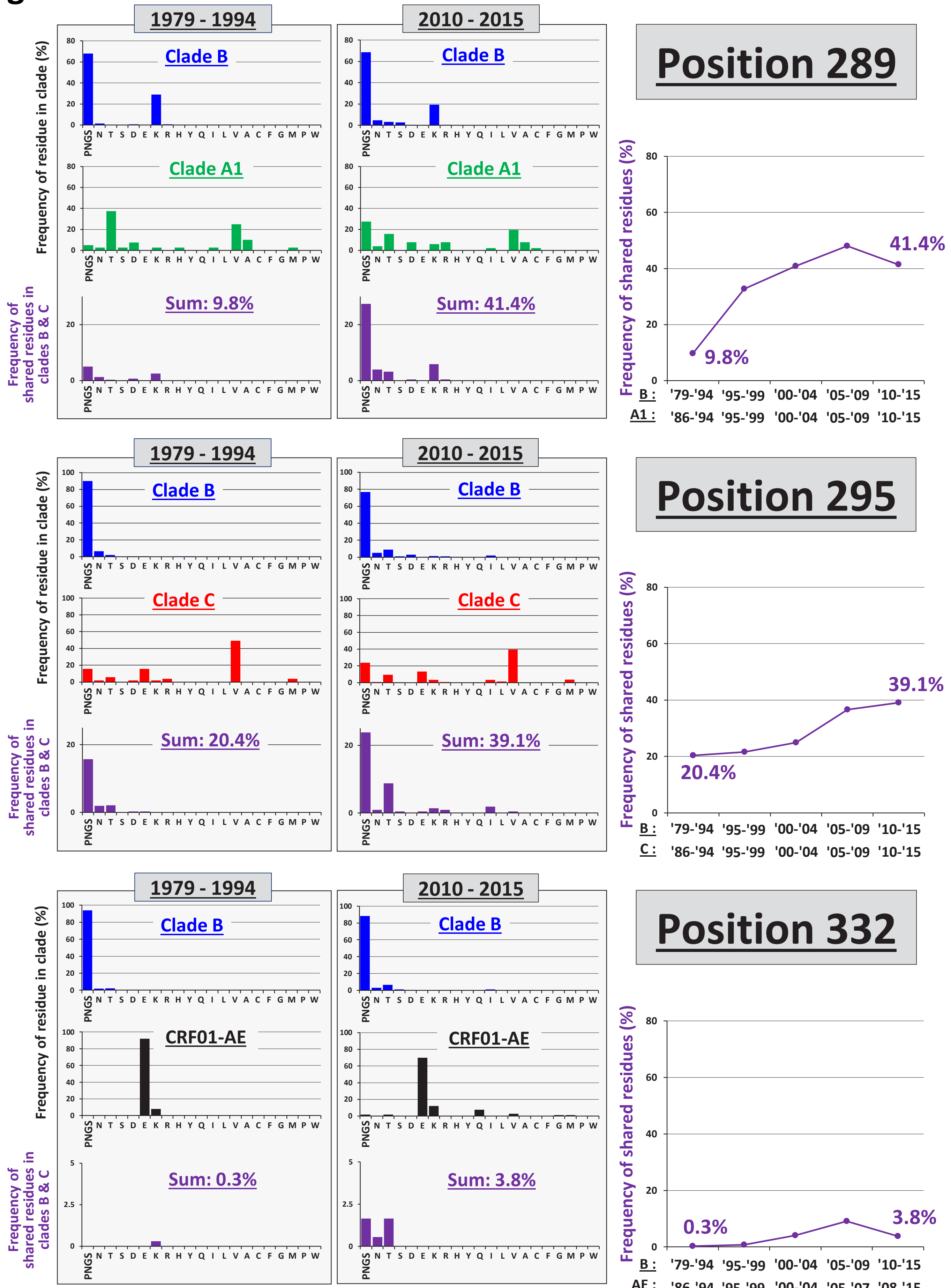
1979 - 1994
2010 - 2015
Position 289
Clade B
Clade A1
Sum: 9.8%
Sum: 41.4%
Frequency of residue in clade (%)
Frequency of shared residues in clades B & C
Frequency of shared residues (%)
9.8%
41.4%
B : '79-'94 '95-'99 '00-'04 '05-'09 '10-'15
A1 : '86-'94 '95-'99 '00-'04 '05-'09 '10-'15
Position 295
Clade C
Sum: 20.4%
Sum: 39.1%
20.4%
39.1%
B : '79-'94 '95-'99 '00-'04 '05-'09 '10-'15
C : '86-'94 '95-'99 '00-'04 '05-'09 '10-'15
Position 332
CRF01-AE
Sum: 0.3%
Sum: 3.8%
0.3%
3.8%
B : '79-'94 '95-'99 '00-'04 '05-'09 '10-'15
AE : '86-'94 '95-'99 '00-'04 '05-'07 '08-'15
PNGS N T S D E K R H Y Q I L V A C F G M P W

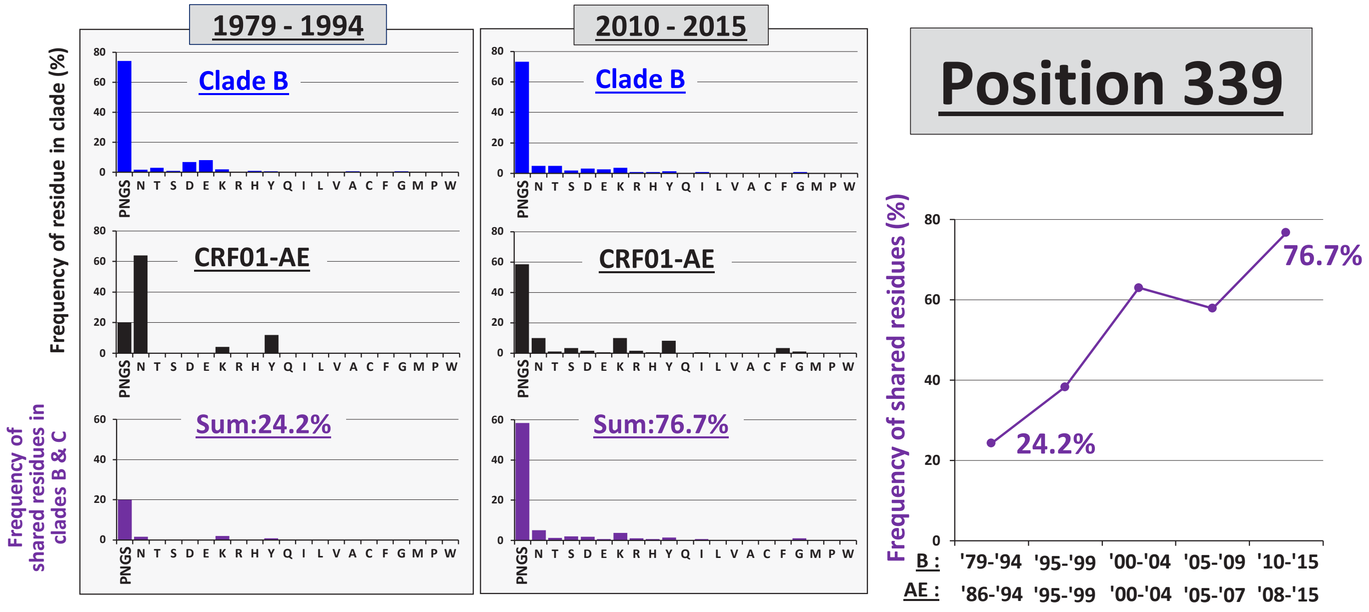
1979 - 1994
2010 - 2015
Position 339
Frequency of residue in clade (%)
Clade B
CRF01-AE
Frequency of shared residues in clades B & C
Sum:24.2%
Sum:76.7%
PNGS N T S D E K R H Y Q I L V A C F G M P W
Frequency of shared residues (%)
24.2%
76.7%
B : '79-'94 '95-'99 '00-'04 '05-'09 '10-'15
AE : '86-'94 '95-'99 '00-'04 '05-'07 '08-'15

### Fig. S8

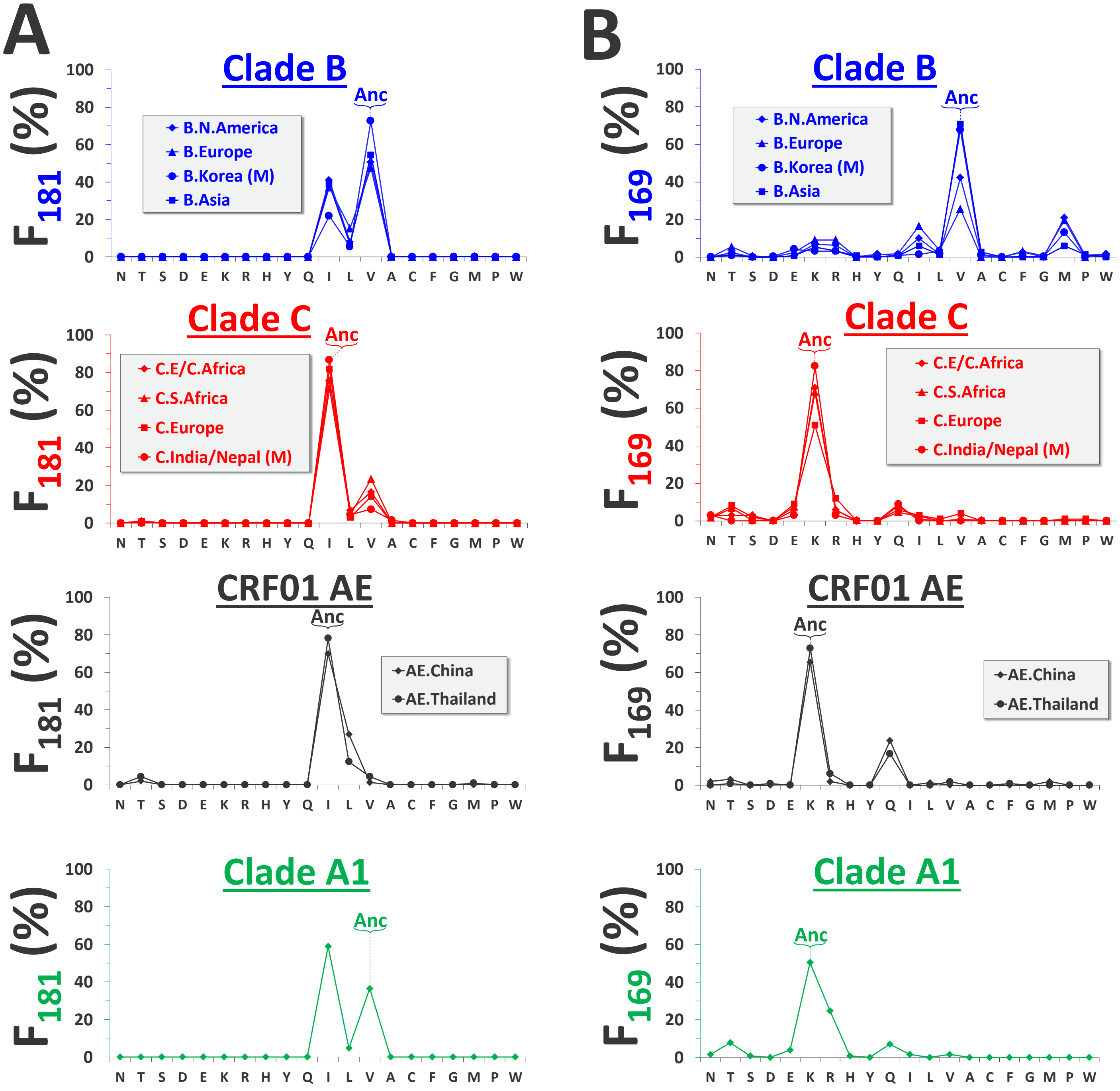
A
B
Clade B
Anc
B.N.America
B.Europe
B.Korea (M)
B.Asia
Clade C
C.E/C.Africa
C.S.Africa
C.Europe
C.India/Nepal (M)
CRF01 AE
AE.China
AE.Thailand
Clade A1
F181 (%)
F169 (%)
N T S D E K R H Y Q I L V A C F G M P W

**Fig. S9**

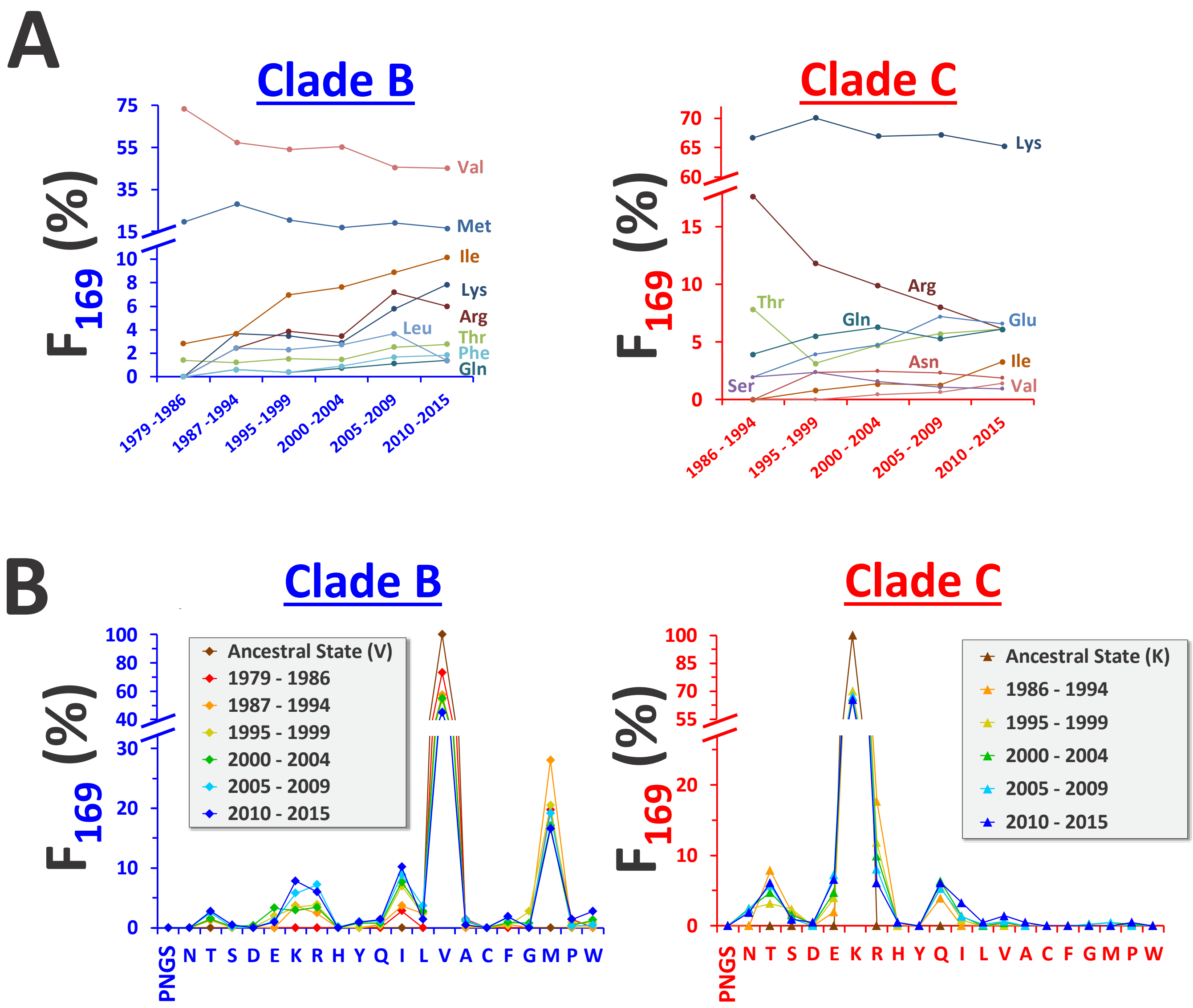
A
Clade B
Clade C
$F_{169}$ (%)
75
55
35
15
10
8
6
4
2
0
Val
Met
Ile
Lys
Arg
Leu
Thr
Phe
Gln
1979 -1986
1987 -1994
1995 -1999
2000 -2004
2005 -2009
2010 -2015
70
65
60
15
10
5
0
Lys
Arg
Thr
Gln
Glu
Asn
Ile
Ser
Val
1986 - 1994
1995 - 1999
2000 - 2004
2005 - 2009
2010 - 2015
B
Clade B
Clade C
Ancestral State (V)
1979 - 1986
1987 - 1994
1995 - 1999
2000 - 2004
2005 - 2009
2010 - 2015
100
80
60
40
30
20
10
0
PNGS N T S D E K R H Y Q I L V A C F G M P W
Ancestral State (K)
1986 - 1994
1995 - 1999
2000 - 2004
2005 - 2009
2010 - 2015
100
85
70
55
20
10
0
PNGS N T S D E K R H Y Q I L V A C F G M P W

### Fig. S10

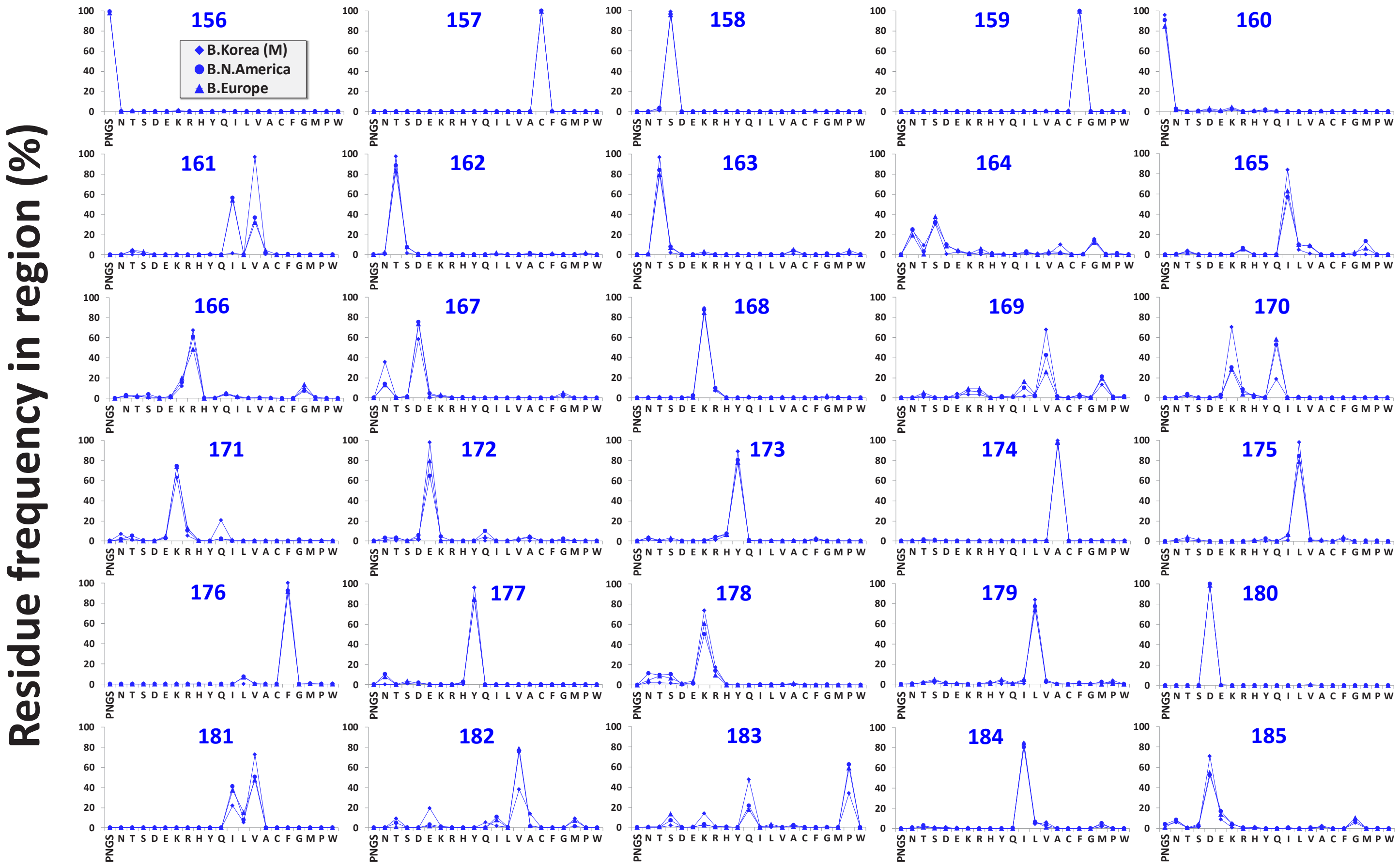
Residue frequency in region (%)
B.Korea (M)
B.N.America
B.Europe
PNGS N T S D E K R H Y Q I L V A C F G M P W

### Table S1

| | Geographic Region | # of Envs[a] | Countries | Tree File | Year range of 'current strain' group[b] | Number of Envs in each time period |
|---|---|---|---|---|---|---|
| **Clade B** (1942 patients) | North America (NA) | 924 | Canada, Cuba, Dominican Republic, Haiti, Jamaica, Trinidad Tobago, United States | Fig S1A | 2007- 2015 (301) | 1979-1986: 71<br>1987-1994: 155<br>1995-1999: 255<br>2000-2004: 535<br>2005-2009: 700<br>2010-2015: 217<br>2016 : 9 |
| | Europe (EU) | 330 | Belgium, Switzerland, Cyprus, Germany, Denmark, Spain, France, Great Britain, Italy, Netherland, Sweden | Fig S1A | 2007- 2015 (162) | |
| | Korea (KR) | 474 M=416 | S. Korea | Fig S1A | 2001- 2009 (312, M=280) | |
| | Asia (AS) | 212 | China, Hong Kong, India, Japan, Myanmar, Philippines, Singapore, Thailand, Taiwan | Fig S1A | 2005- 2015 (151) | |
| **Clade C** (1248 patients) | Southern Africa (SA) | 657 | South Africa and Botswana | Fig S1B | 2007- 2015 (239) | 1986-1994: 51<br>1995-1999: 125<br>2000-2004: 434<br>2005-2009: 464<br>2010-2015: 174 |
| | Eastern/Central Africa (ECA) | 348 | Burundi, Djibouti, Ethiopia, Kenya, Malawi, Tanzania, Somalia, Uganda and Zambia | Fig S1B | 2007- 2015 (103) | |
| | Europe (EU) | 87 | Belgium, Switzerland, Cyprus, Germany, Denmark, Spain, Finland, France, Great Britain, Italy, Netherlands, Portugal, Sweden | Fig S1B | 2005- 2015 (64) | |
| | India/Nepal (IN/NP) | 109 M=94 | India, Nepal | Fig S1B | 2000- 2015 (70, M=61) | |
| **CRF01_AE** (543 patients) | Thailand (TH) | 283 | Thailand | Fig S1C | 2007- 2015 (114) | 1986-1994: 25<br>1995-1999: 62<br>2000-2004: 56<br>2005-2007: 217<br>2008-2015: 183 |
| | China (CN) | 191 | China | | 2007- 2015 (156) | |

| | | | | | | |
|---|---|---|---|---|---|---|
| **Clade A1** (335 patients) | All | 340 | Democratic Republic of Congo, Cameroon, Gambia, Kenya, Niger, Nigeria, Rwanda, Tanzania, Uganda, South Africa, Botswana, Belgium, Canada, Switzerland, Cyprus, Great Britain, Italy, Finland, India, Nepal, Korea, Pakistan, Sweden, United States | Fig S1D | 2006- 2015 (122) | 1985 : 1<br>1986-1994: 39<br>1995-1999: 51<br>2000-2004: 52<br>2005-2009: 145<br>2010-2015: 47 |

#### Table S2

| | | Ancestral Sequence[a] | | | |
|---|---|---|---|---|---|
| Env Region | Position | B | C | 01_AE | A1 |
| C1 | 88 | * | * | * | * |
| V1 | 156 | * | * | * | * |
| V2 | 160 | * | * | * | * |
| | 197 | * | * | * | * |
| C2 | 234 | * | * | * | * |
| | 241 | * | * | * | * |
| | 262 | * | * | * | * |
| | 276 | * | * | * | * |
| | 289 | * | * | * | Thr |
| | 295 | * | Val | * | * |
| V3 | 301 | * | * | * | * |
| C3 | 332 | * | * | Glu | * |
| | 339 | * | * | Asn | * |
| | 356 | * | * | * | * |
| V4 | 386 | * | * | * | * |
| | 392 | * | * | * | * |
| C4 | 448 | * | * | * | * |
